# Multiple plasmid origin-of-transfer substrates enable the spread of natural antimicrobial resistance to human pathogens

**DOI:** 10.1101/2020.04.20.050401

**Authors:** Jan Zrimec

## Abstract

Antimicrobial resistance poses a great danger to humanity, in part due to the widespread horizontal transfer of plasmids via conjugation. Modeling of plasmid transfer is essential to uncovering the fundamentals of resistance transfer and for development of predictive measures to limit the spread of resistance. However, a major limitation in the current understanding of plasmids is the inadequate characterization of the DNA transfer mechanisms, which conceals the actual potential for plasmid transfer in nature. Here, we consider that the plasmid-borne origin-of-transfer substrates encode specific DNA structural properties that can facilitate finding these regions in large datasets, and develop a DNA structure-based alignment procedure for typing the transfer substrates that outperforms mere sequence-based approaches. We identify thousands of yet undiscovered DNA transfer substrates, showing that actual plasmid mobility can in fact be 2-fold higher and span almost 2-fold more host species than is currently known. Over half of all mobile plasmids contain the means to transfer between different mobility groups, which links previously confined host ranges across ecological habitats into a robust plasmid transfer network. We show that this network in fact serves to transfer antimicrobial resistance from the environmental genetic reservoirs to human pathogens, which might be an important driver of the observed rapid resistance development in humans and thus an important point of focus for future prevention measures.

## 1. Introduction

Horizontal gene transfer of antimicrobial resistance (AMR) genes occurs via the processes of transformation and conjugation. The former mediates especially narrow range, intra-genus transfers ^1,2^, whereas the latter is implicated in a wider range of transfer hosts ^3,4^ and potentially enables AMR to overcome the toughest phylogenetic and ecological transmission barriers ^5–10^. Consequently, the interaction between conjugative relaxase enzymes and their DNA origin-of-transfer (oriT) substrates facilitates the majority of all AMR transfers in nature ^11,12^ and is especially important for ones related to human infection complications ^13^. However, the current knowledge on conjugative transfer mechanisms and systems ^4,14–16^ is unable to describe the unprecedented amount of observed horizontal transfer ^7,8,12,17^ that seems to transcend all transfer barriers between resistance reservoirs and human hosts ^5–7,9,10,12,13,18^.

The standard approach for characterization of plasmid mobility involves classification of conjugation and mobilization genes ^14^, especially typing of relaxase enzymes into the respective mobility (Mob) groups ^15,19^. However, besides the possibility of yet unidentified enzymes and mobility groups ^15,20–24^, multiple new processes have recently been uncovered that might confer additional mobility to plasmids and involve the origin-of-transfer (oriT) DNA substrate. These include (i) broadened relaxase binding specificities to multiple different oriT sequence variants ^25–29^, which, according to the evolutionary theory of such DNA regions ^4,30,31^, indicates the possibility of plasmids carrying multiple functional secondary oriTs, and (ii) trans-mobilization of plasmids carrying oriTs triggered by relaxases from co-resident plasmids acting *in trans* on the non-cognate oriTs ^32–35^. The latter mechanism demonstrates that oriT regions are the only elements of the conjugation machinery required in *cis* ^21^ and suggests that many plasmids classified as non-mobile due to the absence of putative relaxases may in fact be mobilizable ^33^. However, although typing the oriT enzymatic substrates instead of the genetic scaffolds might present improvements to the current understanding of plasmid mobility, no systematic studies of oriTs across sequenced plasmids have yet been performed, likely due to the lack of available data and tools that would enable such oriT typing.

A major problem with uncovering oriT regions is that, apart from being experimentally laborious, it is computationally challenging due to multiple molecular mechanisms and a variety of DNA sequence elements present and coevolving in the DNA substrate ^4^, even among plasmids belonging to a single species such as *Staphylococcus aureus* ^32^. OriT typing requires algorithms beyond simple sequence-based alignment ^36,37^, which can recognise and process more complex molecular motifs such as inverted repeats and hairpins ^38–40^ as well as underlying DNA physicochemical and conformational features. These underpin key protein-DNA readout and activity mechanisms ^4,41–43^ as well as define conserved niches of structural variants that enable good resolution between Mob groups and subgroups ^4^. The use of DNA structural representations has indeed led to improvements in algorithms for identification of other regulatory regions, such as promoters and replication origins ^44–48^. Despite this, instead of using tools that probe the actual relaxase-oriT interaction potential by identifying molecular properties that are the basis of such interactions, conventional approaches for oriT analysis still rely on mere nucleotide sequence-based methods ^32,36,37^.

Hence, here we prototype a DNA structure-based alignment algorithm for finding oriT variants, enabling us to locate and Mob-type oriT regions across thousands of sequenced plasmids. Based on the newly uncovered oriT variants, since they can facilitate both *in cis* and *in trans* plasmid transfer, we re-analyse the amount of potential mobile plasmids and mobile plasmid-carrying host species. We then evaluate if and how the uncovered fraction of oriTs helps to overcome the known barriers to horizontal gene transfer, by reconstructing and analysing the network of potential AMR transfers between different species and habitats, especially those from the environmental reservoir to the human flora.

## 2. Results

### Structural alignment algorithm improves oriT-typing performance

Since a DNA alignment algorithm performs multiple comparisons between a query and a target sequence by evaluating a distance function, we first developed a structural distance function, termed s-distance (Fig S1-1, Methods M2). This enabled comparison between structurally encoded oriTs and non-encoded ones, where the ungapped p-distance was used. For the comparison we used a balanced dataset of 64 oriT regions from 4 Mob groups ^4^ (Methods M1), where region sizes were varied stepwise to cover a single relaxase enzymatic site of 40 bp up to the whole oriT region of 220 bp containing multiple binding sites (Fig 1A). Furthermore, the comparison included discrimination of both (i) positive oriTs (aligned to nic site) from negative non-oriT sequences and (ii) Mob groups. With structurally encoded DNA, significant (Anova *p*-value < 1e-4) discrimination of both positive/negative and Mob groups was achieved in the whole oriT size range (Fig 1B,C), compared to the bare nucleotide sequences, where results were significant (Anova *p*-value < 0.05) only with oriT regions equal to or shorter than 120 bp. This was corroborated with the Jaccard distance, which significantly (Ranksum *p*-value < 1e-16) decreased by 40% with increasing oriT region size when using structurally encoded k-mers (Methods M2, Fig S1-2), whereas it increased with nucleotide k-mers (Ranksum *p*-value < 1e-9). The results suggested that our structural encoding approach leveraged the chemical information in longer query regions and could thus improve multiple sequence comparisons with alignments by increasing the statistical depth (Fig S1-3).

**Figure 1.**
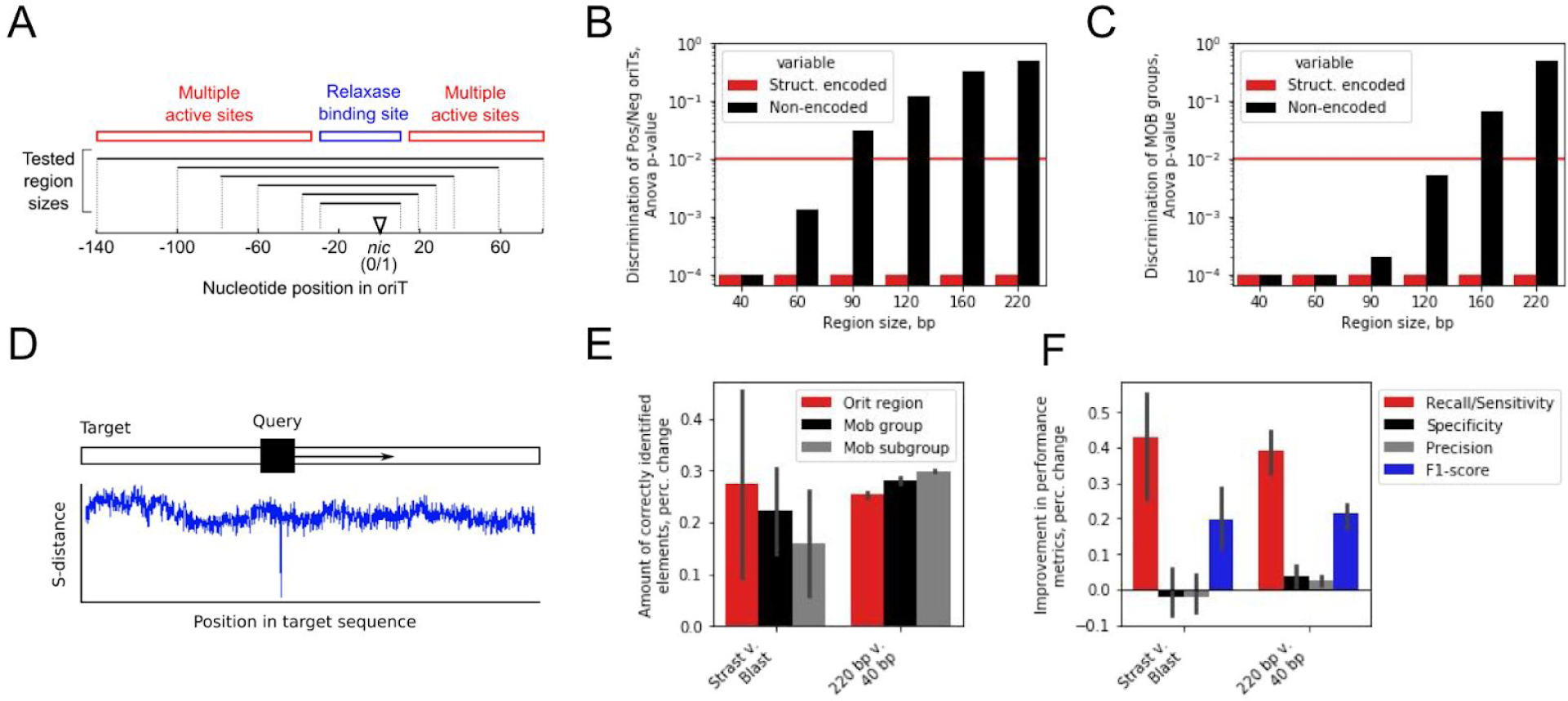
Structural alignment algorithm improves oriT-typing performance. (A) Schematic depiction of the oriT region and different analysed oriT subsets of 40, 60, 90, 120, 160 and 220 bp, which span the single relaxase binding site or multiple protein recognition and binding (ie. ‘active’) sites, respectively. (B, C) Statistical analysis of nucleotide and structural representations (Methods M3) with different oriT subsets for (B) Mob group discrimination and (C) discrimination of positive and negative examples. (D) Schematic depiction of the structural alignment algorithm, which finds the positions in the target dataset with minimum s-distance to the query sequences. (E) Comparison of the amount of correctly identified elements between our algorithm (Strast) and Blast, and by using 220 bp or 40 bp oriT subsets, for oriT typing as well as discrimination of MOB groups and subgroups. Error bars denote 95% confidence intervals. (F) Comparison of machine learning performance metrics between our algorithm (Strast) and Blast, and by using 220 bp or 40 bp oriT subsets. Error bars denote 95% confidence intervals.

We next prototyped an alignment framework (Fig 1D) that employed the s-distance measure to find target hits to query oriTs, where *p*-values were obtained via permutation tests (Fig S1-1, Methods M2). A 4 Mob group dataset of 106 query oriT regions and 170 testing plasmids ^15,36,49^ was compiled (Fig S1-4, Table S1-1, Methods M1) to test the algorithm’s performance by assessing the oriT/nic location and Mob type of the highest scoring alignment hits. By using full-length 220 bp query regions, on average, 25% more significant (Permutation test *p*-value < 1e-14) oriT hits were recovered and Mob group predictions increased 28% compared to using 40 bp query size (Fig 1E, Fig S1-5). This corroborated that the use of longer queries indeed led to improved algorithm performance (Fig 1B,C). Furthermore, compared to Blast ^37^ and to non-encoded DNA using the p-distance (Methods M2), our approach uncovered, on average, 27% more significant (Permutation test *p*-value < 1e-10) oriT hits and correctly predicted 22% more Mob groups (Fig 1E, Fig S1-5). By analysing machine learning metrics to better understand the algorithm’s performance (Methods M3), we observed that a marked ∼40% increase was achieved with *Recall* (Fig 1F) at a relatively constant *Precision* and *Specificity*, which corresponded to recovering a larger amount of the correct oriTs (Fig S1-6). Our method thus correctly located and Mob typed, on average, 55% of oriTs in the cross-validation and testing datasets (Fig S1-7, S1-8). We further validated the capability of the algorithm to identify specifically *nic* sites using a plasmid dataset with experimentally determined oriT regions that was not Mob typed ^36^ (Methods M1). Out of 13 such plasmids with 14 oriT sites, it correctly identified (Permutational test *p*-value < 1e-12) 6 oriT regions in 5 plasmids with 100% sequence identity and aligned to within ±1 bp of the *nic* sites ^49^ (Fig S1-7, Table S1-2). This indicated that the lack of diversity in the query dataset altogether missed certain oriTs in the testing datasets, which was also confirmed by using smaller query datasets that lowered the algorithm’s performance especially for locating oriT regions (Fig S1-9). Nevertheless, despite the limited oriT data availability, the results experimentally verified the algorithm’s capacity for oriT typing.

### OriT typing reveals a two-fold increase in the number of known mobile plasmids

We used the structural alignment algorithm to explore the diversity of oriT regions in natural plasmids. To cover all available oriT regions, we expanded the query dataset to 112 unique oriTs from 6 Mob groups that included, besides oriTs from the major Mob groups F, P, Q and V, also 3.6% and 0.9% of elements from groups C and T, respectively (Fig 2A, Methods M1). The query dataset covered 59 unique host species with the majority (88.4%) from the phyla *Proteobacteria* and *Firmicutes* (Fig S2-1). The target dataset comprised 4602 natural plasmids with Mob groups determined by relaxase amino acid homology analysis ^50^ (Fig 2B). Here, 28.7% of plasmids were mobile (1307 plasmids that contained 1377 distinct relaxases) with the highest represented Mob groups F, P, Q and V (Fig 2A, Fig S2-2). The target dataset contained 893 distinct host species from 22 distinct phyla, with the mobile plasmids harboured by 40% of the distinct species with 88.1% from the phyla *Proteobacteria* and *Firmicutes* (Fig S2-1).

**Figure 2.**
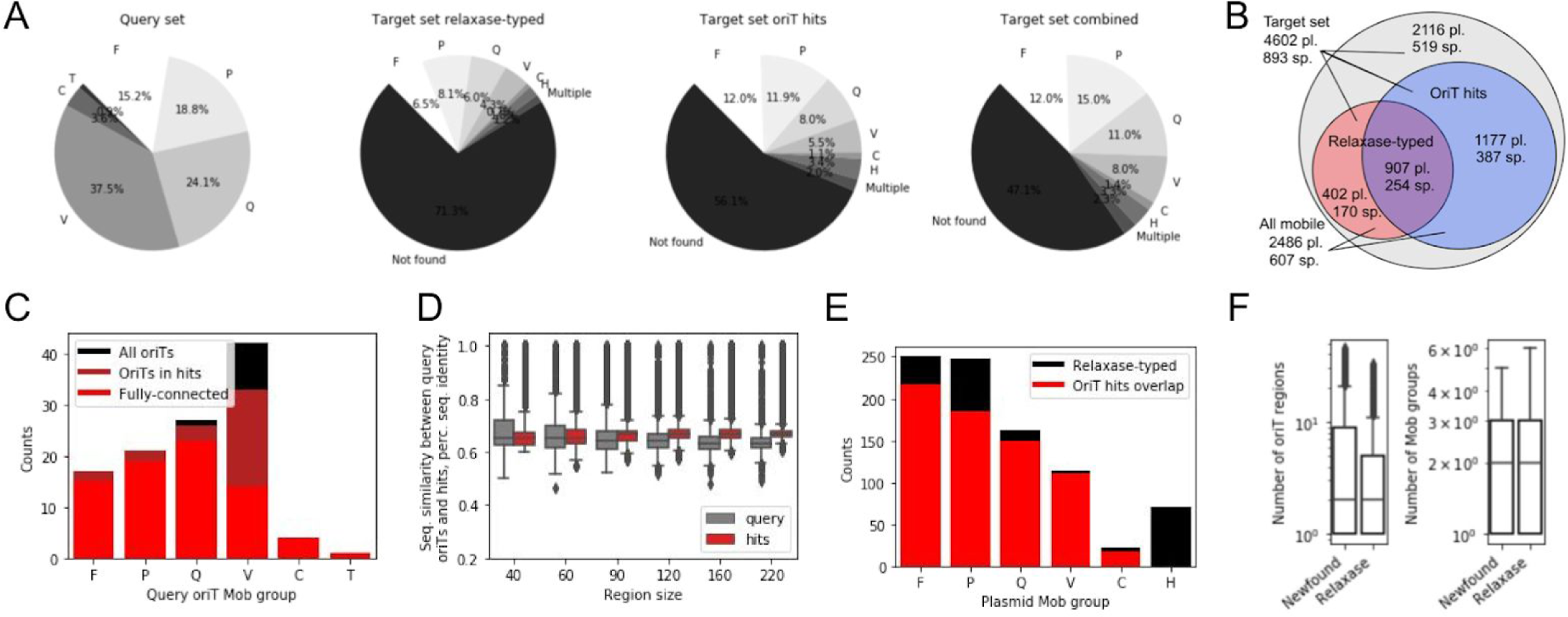
OriT typing reveals a two-fold increase in plasmid mobility. (A) Distribution of Mob groups across the query oriT and target plasmid datasets, where the latter was analysed using either relaxase-typing ^50^, structural alignment-typing or a combination of both methods. (B) Venn diagram of the number of plasmids (pl.) and plasmid-carrying host species (sp.) in the whole target plasmid dataset and the separate subsets uncovered to be mobile by either structural alignment or relaxase-typing. (C) Distribution of Mob groups in the whole query oriT dataset and in the query subsets that returned alignment hits or were present in the fully connected oriT network (see next chapter of Results). (D) Average sequence identities across Mob groups with different oriT size subsets, calculated pairwise between all query oriTs and between each oriT hit and its closest-associated query oriTs within a Mob group. (E) Distribution of Mob groups across relaxase-typed and structural alignment-typed plasmids. (F) Distributions of amounts of oriT regions and Mob groups across the structural alignment-typed (newfound) and relaxase-typed plasmids.

Contrary to the expectations based on the 1377 distinct relaxases found in the target dataset (Fig 2B) that indicated at least a similar amount of oriT regions should be recoverable, with our alignment method we identified 11,497 significant (*q*-value < 1e-8, *E*-value < 0.01) oriT hits (Methods M4). The oriT hits were uncovered with 91% (102/112) of the query regions and covered all Mob groups and subgroups (Fig 2C). They contained sequence features, including sequence homology and inverted repeats (IR) that facilitate relaxase recognition and binding ^4^, similar to those of the query regions and in accordance with published findings showing that relaxases can function with relaxed specificity on non-cognate oriTs with ∼60% sequence homology ^25,26,32^. Indeed, sequence homologies between an oriT hit and its closest-associated query oriT region were above 60% within all the 40 bp relaxase-binding sites and with the majority (> 99.6%) of the larger sized subregions (Fig 2D). Median sequence homologies were well above the ones of the query dataset, where all pairwise seq. homologies within each Mob group were measured (Fig 2D, Methods M4). Furthermore, all the different sizes of oriT subregions were strongly correlated (Pearson’s *r* > 0.730, *p*-value < 1e-16, Table S2-1) between each other as well as to the oriT structural similarity, s-distance (Pearson’s *r* > 0.817, *p*-value < 1e-16, Fig S2-3). In accordance with the analysis of IRs in the query oriTs, where in the 60 bp upstream of *nic* an IR of at least 6 bp could be identified (Fig S2-4A: avg. size was 10 bp, Methods M4), in the oriT hits we found that in fact all but 6 oriTs (0.05%) carried IRs with similar properties (Fig S2-4B: at least 6 bp with an avg. size of 10 bp). Finally, we explored if the uncovered oriTs were located in any specific coding or non-coding regions in the plasmids by obtaining and analysing the CDS records of each plasmid (Methods M3). Indeed, we observed a significant (Fisher’s exact test *p*-value < 1e-16) 4.6-fold increase of oriT presence in non-coding areas and 2-fold decrease in coding ones as well as significant (Fisher’s exact test *p*-value < 1e-16) enrichment in genes related to horizontal mobility, namely conjugation, transposition and integration (Table S2-2).

Out of the 1309 plasmids that carried a relaxase (Fig 2B: 28.4% of target dataset), we identified oriTs in 907 plasmids (69.2% of relaxase-typed plasmids) as well as in an additional 1177 plasmids that were previously not specified as mobile (25.6% of target dataset). Therefore, the uncovered oriT regions combined with the previously typed relaxases resulted in a total of 2486 mobile plasmids (Fig 2B: 54% of target dataset), which represented a 1.9-fold increase in the number of mobile plasmids in the target dataset compared to the initial relaxase typing. Similarly, a 1.7-fold increase in mobile plasmid-carrying host species was identified, when comparing species from the whole set of mobile plasmids (Fig 2B: 607 out of 893 species, 58.2%) with previous relaxase typed ones (356 species, 39.9%). This also corresponded to a 1.4-fold increase in Phyla, with mobile plasmids representing 19 out of the 23 Phyla compared to merely 14 with relaxase typing (Fig S2-1). Furthermore, out of the 907 plasmids where both oriTs and relaxases were identified, the same Mob group, indicating that the oriT was cognate to the relaxase, was identified in 75% of cases (Fig 2E). In the remaining 25% of these plasmids, the uncovered orits could have been secondary oriTs ^30,31^ or corresponded to either unknown or *in trans* acting ^32^ relaxases. The distribution of the oriT-identified Mob groups was found to be comparable to the one expected according to relaxase typing (Fig 2E), whereas a larger diversity of oriTs may exist within certain Mob subgroups than others (Table S2-3). Multiple oriT regions were identified in over 63% of both the relaxase-typed as well as untyped mobile plasmids, where on average, 2 orits from 2 different Mob groups were identified per plasmid (Fig 2F). This supported the notion that, besides secondary and *in trans* oriTs, the untyped plasmids likely carried un-identified relaxases ^15,20–24^.

Since the amount of query oriTs was the limiting factor in our analysis of plasmid mobility, we explored what effect a larger query dataset could have on the findings. Briefly, we simulated the results with a larger query dataset by performing curve fitting on results obtained with 10 repetitions of random 10-fold dilutions of the present dataset (Methods M5). An approximate linear rule was observed between the size of the query dataset and the amount of uncovered oriT-hits, as each order of magnitude increase in oriT hits required likewise an order of magnitude larger query dataset (Fig S2-5A: e.g. 1e5 hits are obtained with a query set of ∼975 oriTs). Consequently, with each order of magnitude increase of the size of the query dataset, approximately 1500 more mobile plasmids (Fig S2-5B: starting from an initial value of 500 with 10 oriT queries) and 250 more mobile plasmid-carrying species were uncovered (Fig S2-5C). Additionally, we observed that in order to achieve a full overlap with the relaxase-typed plasmids, a considerably larger query dataset than is currently available is required, comprising 415 oriTs (95% lower and upper bounds were 328 and 532, respectively, Fig S2-5D). The demonstrated limitations of the query data suggest that the present published results ^50,51^ and our findings might still be an underestimation of the true plasmid mobility present in nature.

### Over one half of mobile plasmids can transfer between different Mob groups

A large part of the newly uncovered oriTs were additional regions to the primary ones that corresponded to the plasmid cognate relaxases (Fig 2F, Fig 3A inset). This resulted in 1331 multi-oriT plasmids (64% of all oriT-typed mobile plasmids, of which 43% were relaxase-typed) that carried on average 5 oriTs (Fig S3-1). First, we analysed the connectivity (co-occurrence network) between the different oriT regions (nodes), when they were shared across the multi-oriT plasmids (edges), which represented an undirected multi-edged graph (Methods M6). Since each oriT hit was characterised only by its closest-associated query oriT, the actual oriT node diversity was limited to the 102 query oriTs that returned hits (Fig 2E). The graph contained a total of 79,004 connections between 552 interacting oriT node pairs (Fig 3B), with an average of 6 connections and up to 3528 connections per oriT node pair (ie. oriTs were shared across this many plasmids, Fig 3A). Among the multiple small connected components in this graph, one was larger than 2 nodes and comprised 76 oriT nodes (75% of all oriTs, Fig 3B) that proportionally represented all 6 Mob groups except Mob V, of which only 38% was in the fully connected subgraph (Fig 2E). The observations of a power law of the degree distribution (Fig S3-2: degree exponent was 0.97) indicated that this oriT network obeyed the laws of natural biological scale-free networks, while the decrease of the clustering coefficient with increasing degree hinted at the possibility of a hierarchical topology (Fig S3-2: global clustering coefficient was 0.47) ^52^. In fact, we observed that specific oriT regions acted as hubs and connected multiple other regions across the Mob groups (Fig 3B). The most highly connected hubs were for instance pNL1-, BNC1 Plasmid 1- and pBBR1-like oriTs from Mob F, Q and V, respectively, that connected over 50 unique oriTs from all 6 Mob groups via ∼1500 connections each (see details in Table S3-1).

**Figure 3.**
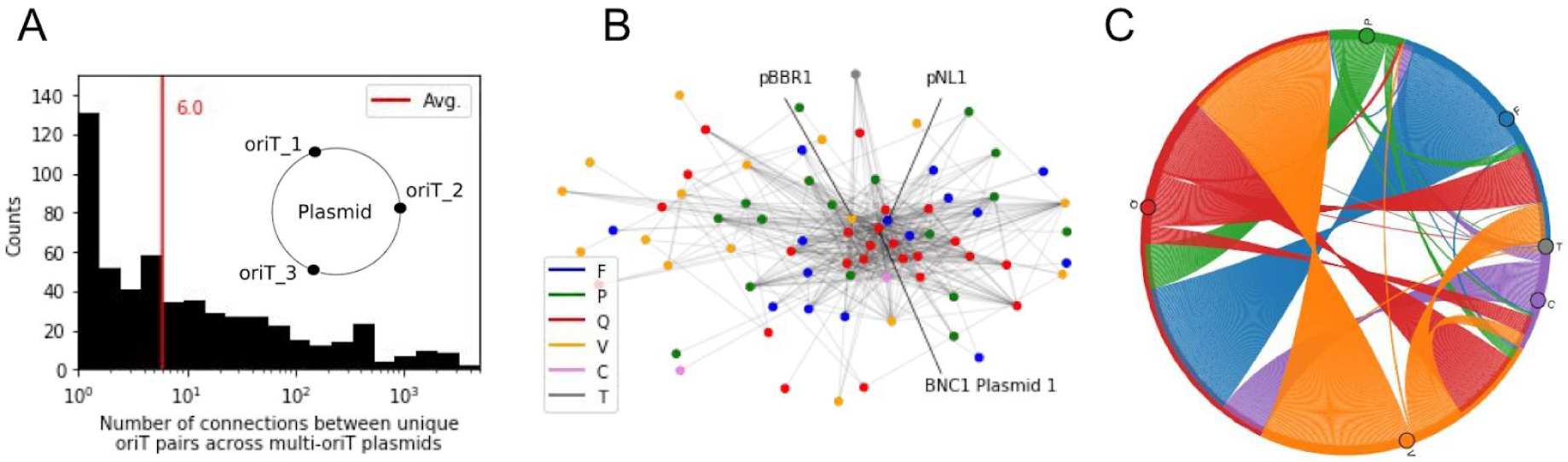
Over one half of mobile plasmids can transfer between different Mob groups. (A) Distribution of the number of shared oriTs across the multi-oriT plasmids, based on similarity to their closest-associated query oriTs and considered here as connectivity between plasmids. Inset: schematic depicting multi-oriT plasmids. (B) The undirected multi-edged graph of shared oriTs (nodes) across plasmids (edges). The most highly connected (co-occurring) oriTs from plasmids pBBR1, pNL1 and BNC1 Plasmid 1 are marked (note: plasmids of closest-associated query oriTs). Colors denote Mob groups. (C) Potential connectivity between Mob groups based on shared oriTs in multi-oriT plasmids (note: undirected graph).

We next focused on the connectivity of Mob groups, since over 90% of the multi-oriT plasmids contained on average 2 unique Mob groups and 3 unique Mob subgroups (Fig S3-1). Indeed, the number of different identified Mob groups per plasmid increased proportionally to the amount of oriT regions (Pearson’s *r* = 0.799, *p*-value < 1e-16). Compared to relaxase-typing, we observed on average a 7-fold increase of identified oriT elements per Mob group, which led to a 75-fold increase in the amount of connections, with the highest increases in Mob groups F, Q, and C (above 75-fold increase) and lowest in Mob groups P and V (11 and 13-fold increases, respectively, Table S3-2). Therefore, by switching the nodes of the above oriT graph with the corresponding Mob groups, we were able to analyse the high connectivity between Mob groups within the plasmids (Fig 3C, Table S3-3, Methods M6). In correspondence with the most frequently co-occurring Mob groups F, Q and V (Fig S3-2), the highest flux was also observed between these groups. The most strongly connected and highly promiscuous was Mob Q, as 35,062 connections were measured within Mob Q, 15,081 connections between Q and F, and 12,637 connections between Q and V (Table S3-3). Due to the large amount of intra-Mob fluxes, we also analysed the connectivity of Mob subgroups. This showed that the largest amounts of connections frequently involved the unknown subgroups, specifically Mob Qu with Q2, Fu, V2 and P7 (Table S3-4), pointing to possible insufficient typing and underestimation of the true mobility within the Mob groups. Overall, the results demonstrate how each multi-oriT plasmid contains the initial means of transfer between multiple Mob subgroups and groups, and can thereby in principle connect multiple different plasmid host species either transiently, such as via transfer hosts ^53,54^, or via replicative hosts ^3,4^.

### High connectivity across habitats facilitates transfer of resistance genes to humans

We explored how the increased potential for plasmid transfer between different host species (Fig 3) contributes to the connectivity across different environmental habitats and whether it facilitates the spread of AMR genes to humans. For this task, we expanded the plasmid and oriT data by mapping the host species across 9 habitat supertypes (Table S4-1) according to published data ^55–60^ (Methods M6). This retained 43% (227 of 532) of the unique species carrying plasmids, where multiple oriTs and habitat sizes reflected those of the full habitat dataset (according to the amount of unique species) but were on average 8-fold smaller (Fig S4-1). Next, we constructed a directed graph representation of habitat nodes connected by potential plasmid transfers as edges, where habitats of donor hosts carrying the mobile plasmids (outbound connections) connected to habitats of potential acceptor hosts deduced from the query oriTs (inbound connections, Fig 4A, Fig S4-2, Methods M6). The network comprised 141,395 connected habitat node pairs, comprising 1,600,978 plasmid connections between the habitats (Fig 4A). On average, each plasmid was present in 6 habitats and had access to 5 unique host species (Fig 4B) as well as 6 habitats (Fig S4-3). The number of mobile plasmids and oriTs across the habitats were both strongly correlated to the number of unique species (Pearson’s *r* was 0.961 and 0.720, *p*-value < 0.03, respectively), with an average of 3 oriTs per plasmid (Fig S4-4 and S4-5). Interestingly, among the highest amount of oriTs per plasmid was found in industrial (includes food production and water treatment facilities) and animal habitats, known to harbor resistance ^61^ (Fig S4-4).

**Figure 4.**
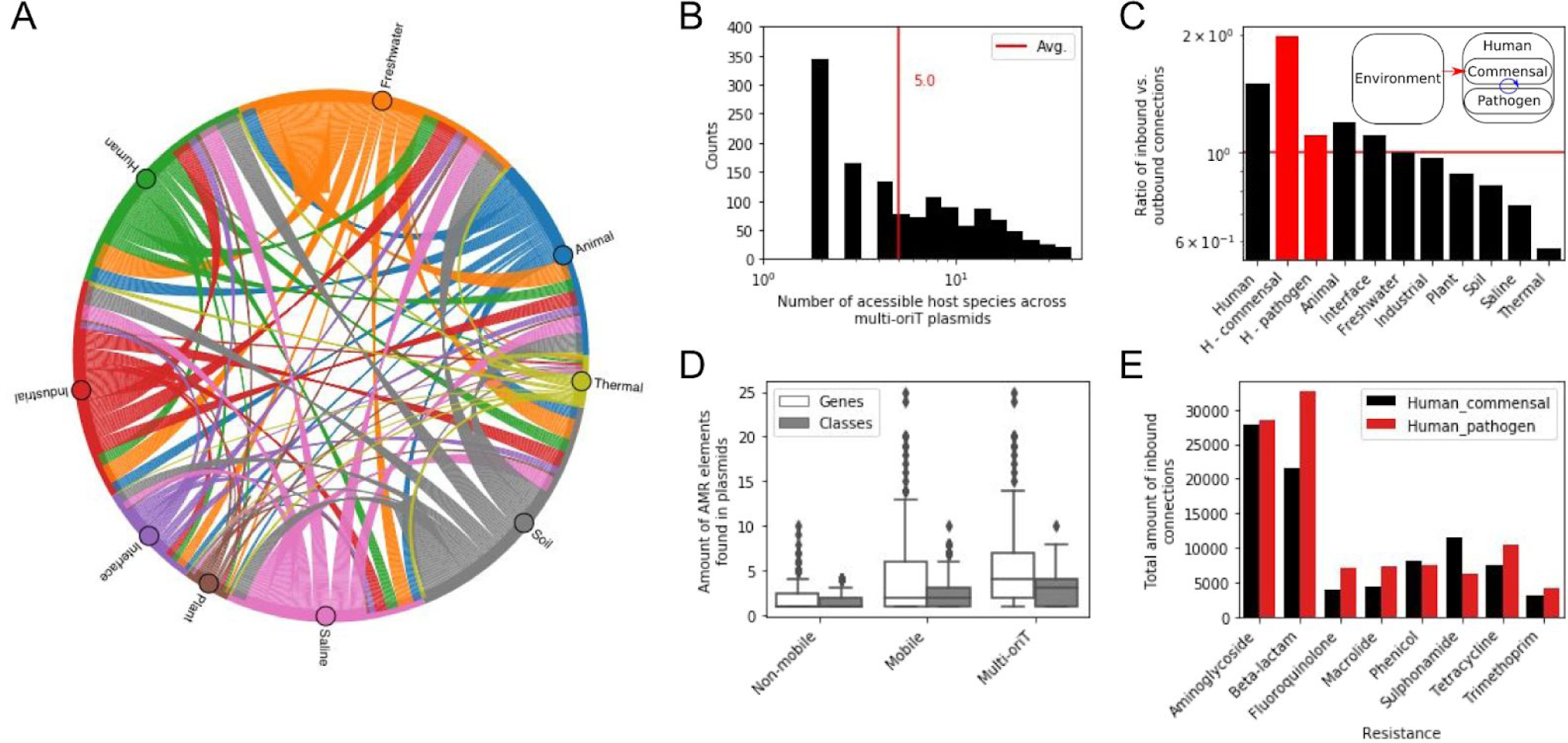
High connectivity across habitats facilitates transfer of resistance genes to humans. (A) Directed graph depicting the connectivity of habitats (nodes) based on potential plasmid transfers (edges). Outbound connections are based on habitats of donor hosts carrying the mobile plasmids and inbound connections are habitats of potential acceptor hosts deduced from the query oriTs. (B) Distribution of the number of host species accessible to the multi-oriT plasmids. (C) Ratio of inbound vs. outbound connections across different habitats, where Human commensal and pathogen microbiomes are highlighted in red. Inset depicts how the Human commensal flora acts as a buffer for transfer of AMR from environmental reservoirs the Human pathogenic flora. (D) Amount of AMR genes and classes found in non-mobile and mobile plasmids. (E) Amount of inbound connections of different AMR classes to the Human microbiome. Only AMR classes with more than 2000 connections shown (see Fig S4-9).

The number of outbound and inbound connections differed across the habitat types (Fig S4-6), as they were significantly correlated with the number of unique species (Pearson’s *r* was 0.895 and 0.916, *p*-value < 2e-3, respectively, Fig S4-5). However, despite the strong correlation (Pearson’s *r* = 0.894, *p*-value < 2e-3) between outbound and inbound connections across habitats, we observed a marked increase in the ratio of inbound vs. outbound connections in the Human and Animal habitats compared to the others (Fig 4C: 61% and 29% increase, respectively). Conversely, the Thermal habitat displayed 43% less inbound than outbound connections, which might also have been due to the lack of representation of plasmids from that habitat ^55^ or the lowest amount of species diversity (Fig S4-1). In order to analyse the transfers to and within the Human habitat more comprehensively, we expanded the habitat taxonomy to include commensal and pathogen types as well as tissue subtypes (Table S4-1, Methods M6). Strikingly, a 2-fold higher amount of inbound vs. outbound connections was observed with commensals (Fig 4C), whereas pathogens displayed a mere 11% increase. The highest amounts of connections to humans were from the Freshwater and Soil habitats (Fig S4-7A), especially via the gut and oral microbiomes (Fig S4-8A). Notably, all tissues except respiratory ones displayed an increase of inbound connections compared to outbound ones, with unassigned bacteria harboring some of the most highly receptive species (Fig S4-8B). Further analysis of the transfer within the human system itself showed an approximately equal rate of outbound and inbound connections (Fig S4-7B), suggesting that commensal bacteria act as the main interface for horizontal uptake of genes from the environment ^62^, which they then further disseminate to the pathogens within the human body (Fig 4C inset).

By predicting the AMR genes in the plasmids (Methods M6), we found a moderate positive correlation (Pearson’s *r* = 0.462, *p*-value < 1e-16), between the number of oriTs and the number of resistance genes in a plasmid (Fig 4D). Thus, 33% of the multi-oriT plasmids carried on average 4 genes from 3 different AMR classes (Fig 4D). When viewed from the perspective of the above cross-habitat transfer network (Fig 4A), the amount of inbound connections of resistance genes from the environment to the human flora indicated that, as expected, the most abundant flow of genes corresponded to the oldest and most widely used classes of antibiotics, for which also resistance is most developed and widespread ^63^ (Fig 4A, Fig S4-8). Moreover, in this case, the amount of inbound connections to pathogens surpassed that of commensals by almost 20% (Fig 4E), supporting that AMR transfer routes generally serve a different portion of microbes compared to plasmid transfer in general (Fig 4C).

## 3. Discussion

Here, we explore the potential for horizontal transfer of natural plasmids by attempting to identify all conjugative DNA origin-of-transfer substrates coded within them. By prototyping a structural alignment approach to find and characterize oriT regions across plasmids (Fig 1D), we discover an almost 8-fold larger amount of oriTs than expected according to relaxase typing (Fig 2B). Analysis of these regions suggests that the actual number of transferable plasmids can be as much as 2-fold higher and span almost 2-fold more host species than is currently known (Fig 2B), with the amount of plasmid-borne AMR genes in fact proportional to this assessment of plasmid mobility (Fig 4D). We show that over half of all mobile plasmids contain the means to transfer between different mobility groups (Fig 3C), potentially linking multiple host ranges that were previously thought to be confined ^3,4^. By analysing this network of potential plasmid transfers across ecological habitats (Fig 4A), we find that human and animal microbiomes display a large amount of plasmid connections with the environment. The considerably larger influx of plasmids to humans and animals compared to other environmental habitats (Fig 4C) might be a consequence of the increased amount of AMR transfers to these organisms ^63^. In accordance with previous findings ^62^, the network shows that human commensals might act as the main interface for horizontal uptake of genes from the environment in general (Fig 4C), whereas, unsurprisingly, the transfer of the specific widespread AMR genes is generally more highly targeted at pathogens (Fig 4E). Nevertheless, the potential increase in plasmid mobility uncovered here might be an important driver of the observed rapid resistance development in humans ^63,64^ and thus an important point of focus for future prevention measures.

Our oriT-typing procedure is a result of rationally expanding DNA alignment algorithms to incorporate enzymatically-relevant properties of the oriT substrates (Fig 1A), where the conservation of structural properties is detected across the whole 220 bp region compared to mere ∼40 bp of nucleotide sequence in the core relaxase-binding site ^4^ (Fig 1C,D). By allowing the use of at least 2-fold longer query sequences, structural alignment achieves a much larger statistical depth than sequence alignment (Fig S1-3), which means that oriTs can be efficiently sought across whole plasmids instead of just the vicinity of relaxases, as is the constraint with current methods ^36^ (Fig 1E). Since, due to the nature of the conserved structural properties, each enzymatic substrate corresponds to multiple possible sequence variants, the benefit of the DNA structural encoding is that it exposes these sequence variants by accessing the search space of the enzymatic co-evolutionary constraints (i.e. DNA structural background) ^4^ (Fig 1A and 2D). The identified candidate oriT regions serve as starting points that can be further verified by typing other known molecular features ^32^, such as inverted repeats ^65,66^ (Fig S2-4) and nucleotide sequence properties of the core enzymatic binding ^38,67^ and nicking sites ^49,68^ (Fig 2D).

Besides finding the majority of expected oriT regions of known cognate relaxases (Figs 2E), almost ⅔ of the oriT-bearing plasmids in fact carry multiple oriTs (Fig 2F). The newly discovered oriTs are frequently located where they are expected, in non-coding regions and within genes related to horizontal mobility (conjugation, transposition and integration) ^69^. However, the amount of Mob groups and depth of enzymatic substrate diversity that could be analysed within each group was constrained by the size of the query set of available oriT regions and nic sites ^19^ (Fig S1-9 and S2-5). By simulating the availability of a larger set of query sequences, a linear relationship between the amount of uncovered and query oriTs indicates that our current sampling is likely still an underestimation of the actual plasmid transfer potential that could possibly span all plasmids ^51,70^ (Fig 2I). Our results of a 2-fold higher plasmid mobility compared to relaxase typing, with an almost similar increase in the amount of mobile plasmid-bearing host species (Fig 2B), point to multiple possibilities that further undermine the paradigm of a one relaxase-one oriT conjugative plasmid system spanning less than ⅓ of plasmids ^50^: (i) a massive underidentification of relaxase enzymes ^51,71^, (ii) relaxase promiscuity ^21,31^ and oriT evolutionary mechanisms ^30^ leading to many functional secondary oriTs, and (iii) a system-wide adoption of relaxase *in trans* mechanisms ^21,72^.

Since plasmids are vehicles for transfer and long-term storage of ‘common goods’ that include, besides AMR, also virulence, heavy metal resistance and other genes ^73^, one can expect the tendency of the global plasmid transfer network to possess: (i) the capacity of at least some elements to bypass key transfer barriers, including phylogenetic ^2,74^, host range (defined by Mob and Inc/Rep groups, respectively) ^3,4,50^ as well as ecological habitat constraints ^2,75^, and (ii) a robust fault tolerant system that increases the guarantee for transfer as well as information storage ^70,76^. Firstly, we found that the plasmids bearing multiple oriTs connect different Mob groups and subgroups (Fig 3C), thus transcending phylogenetic and host range barriers as well as potentially creating a global cross-habitat network that is reminiscent of a dense information network ^70,77^ (Fig 4A). This network interconnects environmental genetic reservoirs and carriers as well as commensal vectors ^62^, which likely act as long-term and shorter-term storage units ^75^, respectively, with known human pathogens ^64^ (Fig 4C). Secondly, with a glimpse of the oriT network topology via the closest-associated query oriTs reminiscent of scale free and even hierarchical networks (Fig 3B and S3-2), the transfer networks display robust fault tolerant properties. As sparsely connected nodes without many direct neighbors are linked to highly connected hubs, even in case of absence of a large number of nodes, the remaining ones are likely still well connected ^52,78^. The finding that specific types of oriTs serve as hubs linking to most of the other oriT types (Fig 3B,C) suggests that certain types of conjugative transfer mechanisms and their corresponding hosts act as transfer hubs that can bypass horizontal gene transfer barriers (Fig 4A) to ensure the flow of genetic information among the different global microbiomes ^64,79^.

## 4. Methods

### M1. Datasets used for alignments

The full query dataset comprised 112 distinct oriTs from 118 plasmids (merged below sequence similarity of 15%) and 6 Mob groups {F,P,Q,V,C,T} (26 Mob subgroups) with known *nic* sites (Figure S1-1A, Dataset S1). The dataset included (i) 48 experimentally verified oriT regions, of which 34 contained experimentally verified nicking sites and 14 contained putative nicking sites, and (ii) 59 oriT regions with computationally predicted nicking sites. For the initial development and testing of the structural alignment algorithm, the following query and testing datasets were used (Table S1-1). Due to the lack of a sufficient number of elements from Mob groups C, H and T for correct testing (Figure S1-4: below 10 elements per group), a 4 Mob group {F,P,Q,V} version of the query dataset with 106 elements was used. The balanced dataset from 4 Mob groups {F,P,Q,V} used for s-distance testing was a subset of the query data containing approx. 16 elements from each Mob group ^4^. A set of negative examples was obtained for each element by extracting sequences from the neighboring vicinity of oriTs. Specifically, the negative examples were selected randomly from a region 200 to 800 bp upstream and downstream from experimental nic sites (defined by size of smallest plasmid), thus containing different non-orit coding and non-coding regions with low sequence similarity (p-distance > 0.6). The testing dataset of 170 plasmids ^15,36,49^ was compiled from 3 separate datasets that included: (i) 106 plasmid sequences of the 4 Mob query dataset, (ii) 51 plasmids with known oriT locations and Mob sites but unknown nic sites and (iii) 13 plasmids with 14 experimentally determined nic/oriT sites but unknown Mob groups, obtained from the OriTFinder database ^36^.

### M2. Development and testing of alignment algorithms

We developed and tested a DNA structure based alignment algorithm, termed Strast, based on (i) encoding DNA into a structural representation ^4^ and (ii) a procedure that finds the most similar segments of target sequences to query sequences using a structural distance measure. In the first stage of Strast, for each DNA sequence, its set of kmers is transformed into a precalculated structural encoding, termed s-mers ^4^ (Fig S1-1B). The s-mers are based on clustered principal components of values obtained from 64 models of physicochemical and conformational DNA properties (Table S1-3). Here, the s-mer size was set to 7 bp, the number of principal components was 18 (out of 64) to capture over 0.99 of data variance, and the k-means clustering algorithm with k = 128 clusters was used. The s-distance between two DNA sequences is defined as the sum of Euclidean distances between each consecutive pair of s-mers in the DNA sequence encodings (Fig S1-1B). The Jaccard distance between structurally encoded DNA sequences is obtained by using s-mers instead of k-mers. The second stage of Strast queries a target sequence for regions with the most similar and statistically significant s-distance (Fig S1-1C: see algorithm pseudocode, Methods M3).

For comparison of Strast to traditional non-encoded sequence based alignments two algorithms were used: (i) Blast v2.24 ^37^ (www.ncbi.com) and (ii) the Strast procedure using the p-distance measure instead of s-distance. To assess the performance of the alignment algorithms for oriT-typing in target sequences, we evaluated the correctness of both (i) oriT and nic location finding to within ± 1 bp ^49^ and (ii) typing of Mob groups and subgroups. Additionally, the effect of the length of query regions in both types of tests was assessed (Fig 1B).

### M3. Statistical analysis and machine learning metrics

The F-test was performed using Permanova ^80^ with sequence bootstraps. The statistical significance of s-distance scores was evaluated using permutational tests, where bootstrap resampling (n_bootstraps = 1e6 per sequence) of randomly selected query oriT sequences (n_seq = 10) was used to estimate the s-distance scores at different p-value cutoffs (from 1e-6 to 1e-1). Next, to obtain a mapping function of s-distance to permutational p-values in the whole range of 1e-132 to 1e-1 (Fig S1-1D), least squares curve fitting to a second order polynomial function was performed, where the theoretical limit of ∼1e-132 was set to correspond to an s-distance of 0. For additional statistical hypothesis testing, the Python package Scipy v1.1.0 was used with default settings.

The following machine learning performance metrics were used to assess alignment algorithm performance: Precision, Recall/Sensitivity, Specificity, Accuracy, F1-score and Matthews correlation coefficient (Table S1-4). To calculate these metrics, true and false positive and negative counts were obtained from the alignment tests (Methods M2) by considering only the most significant hit per alignment. A true or false positive value was assigned if the result was above a specified significance cutoff and corresponded or did not correspond, respectively, to the known value (nic location, Mob group or subgroup), and alternatively, a false or true negative value was assigned to results below the significance cutoff that corresponded or did not correspond, respectively, to the known value.

### M4. Analysis of alignment hits

The newly uncovered regions were analysed by comparing the features of the oriT alignment hits with those of the query dataset, which included sequence properties and inverted repeats. Sequence homology analysis involved (i) calculation of the sequence homologies of the oriT query dataset within each Mob group (ii) calculation of sequence homologies between each oriT hit and its closest-associated query oriT region, (iii) comparison of the sequence homologies of the oriT query and alignment hit datasets, across the different sized oriT subregions. OriT hits with sequences of their relaxase-binding 40 bp subregions that deviated below 60% seq. homology from their query counterparts were removed. Sequence homology was calculated with the *ratio* function (python-Levenshtein package v0.12), where it equalled the Levenshtein (edit) distance divided by the length of the sequence. Similarly, analysis of the inverted repeats (IRs) involved computation of imperfect IRs in both the oriT query and alignment hit datasets, and oriT hits lacking IRs similar to those in the query set were removed. The Matlab package detectIR v2016-01-19 ^81^ was used with IR size limits of (6, 15) bp and containing at most 2 mismatches. From the initially identified 20,255 oriT hits, 11,497 (57%) were retained (Dataset S2).

### M5. Simulations of plasmid mobility

To estimate the results that would be obtained with a larger oriT query dataset, the following procedure was applied. The oriT alignment results with the 4602 target plasmid dataset were diluted according to 10-fold dilutions of the 102 query regions (Fig 2C) used to identify the hits (10 repetitions were used). Least-squares curve fitting was performed (Python package Scipy v1.1.0) using a linear function and the dataset dilutions - specifically between the size of the query oriT dataset and the variables corresponding to the numbers of oriT hits, mobile plasmids, mobile plasmid carrying species and overlap with relaxase-typed plasmids.

### M6. Analysis of transfer between environmental microbiomes

Host species of the oriT alignment results with the subset of multi-oriT plasmids were mapped across 9 habitat supertypes (see habitat groupings in Tables S4-1, S4-2) according to published data on environmental ^55^ and human microbiomes ^57–60^. The habitat taxonomy was further expanded to include commensal and pathogen types ^57^ as well as human tissue subtypes ^55^. For typing antimicrobial resistance genes in the plasmids, the webserver version of ResFinder v3.2 ^82^ was used with default settings.

To study the connectivity between the different oriT regions or Mob groups as nodes, shared across the multi-oriT plasmids as edges, an undirected multi-edged graph was constructed. To study the connectivity and AMR transfer potential between different habitats, a directed graph was constructed, where habitat nodes were connected by multi-oriT plasmids as edges. Habitats of donor hosts carrying the mobile plasmids (outbound connections) were connected to habitats of potential acceptor hosts, deduced from the query oriTs (inbound connections). The Python package NetworkX v2.2 was used.

### M7. Software

Matlab v2018 (www.mathworks.com) and Python v3.6 (www.python.org) were used.

## Acknowledgements

This work was supported by the Slovenian Research Agency under grant agreement n° [Z2-7257] and was in part carried out at the Faculty of Health Sciences, University of Primorska, Izola, Slovenia. I kindly thank Maria Pilar Garcillán-Barcia, Fernando de la Cruz (UNICAN, Spain) and Joshua Ramsey (Curtin Univ., Australia) for sharing and discussing data, Filip Buric (Chalmers Univ. of Tech., Sweden) for commenting on the manuscript, as well as Tomaz Pisanski (UP-FAMNIT, Slovenia) and Ales Lapanje (IJS, Slovenia) for helpful discussions in their respective fields of research.

## References

1. Gibson, M. K., Forsberg, K. J. & Dantas, G. Improved annotation of antibiotic resistance determinants reveals microbial resistomes cluster by ecology. ISME J. 9, 207–216 (2015).

2. Hu, Y. et al. The Bacterial Mobile Resistome Transfer Network Connecting the Animal and Human Microbiomes. Appl. Environ. Microbiol. 82, 6672–6681 (2016).

3. Garcillán-Barcia, M. P., Alvarado, A. & de la Cruz, F. Identification of bacterial plasmids based on mobility and plasmid population biology. FEMS Microbiol. Rev. 35, 936–956 (2011).

4. Zrimec, J. & Lapanje, A. DNA structure at the plasmid origin-of-transfer indicates its potential transfer range. Sci. Rep. 8, 1820 (2018).

5. Dolejska, M. & Papagiannitsis, C. C. Plasmid-mediated resistance is going wild. Plasmid 99, 99–111 (2018).

6. Ben Maamar, S. et al. Mobilizable antibiotic resistance genes are present in dust microbial communities. PLoS Pathog. 16, e1008211 (2020).

7. Sun, D., Jeannot, K., Xiao, Y. & Knapp, C. W. Editorial: Horizontal Gene Transfer Mediated Bacterial Antibiotic Resistance. Frontiers in Microbiology vol. 10 (2019).

8. Mathers, A. J., Peirano, G. & Pitout, J. D. D. The role of epidemic resistance plasmids and international high-risk clones in the spread of multidrug-resistant Enterobacteriaceae. Clin. Microbiol. Rev. 28, 565–591 (2015).

9. Malhotra-Kumar, S. et al. Colistin resistance gene mcr-1 harboured on a multidrug resistant plasmid. The Lancet infectious diseases vol. 16 283–284 (2016).

10. Wang, B. & Sun, D. Detection of NDM-1 carbapenemase-producing Acinetobacter calcoaceticus and Acinetobacter junii in environmental samples from livestock farms. J. Antimicrob. Chemother. 70, 611–613 (2015).

11. Alekshun, M. N. & Levy, S. B. Molecular mechanisms of antibacterial multidrug resistance. Cell 128, 1037–1050 (2007).

12. von Wintersdorff, C. J. H. et al. Dissemination of Antimicrobial Resistance in Microbial Ecosystems through Horizontal Gene Transfer. Front. Microbiol. 7, 173 (2016).

13. San Millan, A. Evolution of Plasmid-Mediated Antibiotic Resistance in the Clinical Context. Trends Microbiol. 26, 978–985 (2018).

14. Smillie, C., Garcillán-Barcia, M. P., Francia, M. V., Rocha, E. P. C. & de la Cruz, F. Mobility of plasmids. Microbiol. Mol. Biol. Rev. 74, 434–452 (2010).

15. Garcillán-Barcia, M. P., Francia, M. V. & de la Cruz, F. The diversity of conjugative relaxases and its application in plasmid classification. FEMS Microbiol. Rev. 33, 657–687 (2009).

16. Fernandez-Lopez, R., Redondo, S., Garcillan-Barcia, M. P. & de la Cruz, F. Towards a taxonomy of conjugative plasmids. Curr. Opin. Microbiol. 38, 106–113 (2017).

17. Lopatkin, A. J. et al. Persistence and reversal of plasmid-mediated antibiotic resistance. Nat. Commun. 8, 1689 (2017).

18. Salyers, A. A. & Amábile-Cuevas, C. F. Why are antibiotic resistance genes so resistant to elimination? Antimicrob. Agents Chemother. 41, 2321–2325 (1997).

19. Garcillán-Barcia, M. P., Pilar Garcillán-Barcia, M., Redondo-Salvo, S., Vielva, L. & de la Cruz, F. MOBscan: Automated Annotation of MOB Relaxases. Horizontal Gene Transfer 295–308 (2020) doi: 10.1007/978-1-4939-9877-7_21.

20. Ramachandran, G. et al. Discovery of a new family of relaxases in Firmicutes bacteria. PLoS Genet. 13, e1006586 (2017).

21. Guzmán-Herrador, D. L. & Llosa, M. The secret life of conjugative relaxases. Plasmid 104, 102415 (2019).

22. Soler, N. et al. Characterization of a relaxase belonging to the MOBT family, a widespread family in Firmicutes mediating the transfer of ICEs. Mob. DNA 10, 18 (2019).

23. Wisniewski, J. A. et al. TcpM: a novel relaxase that mediates transfer of large conjugative plasmids from Clostridium perfringens. Mol. Microbiol. 99, 884–896 (2016).

24. Coluzzi, C. et al. A Glimpse into the World of Integrative and Mobilizable Elements in Streptococci Reveals an Unexpected Diversity and Novel Families of Mobilization Proteins. Frontiers in Microbiology vol. 8 (2017).

25. Kishida, K., Inoue, K., Ohtsubo, Y., Nagata, Y. & Tsuda, M. Host Range of the Conjugative Transfer System of IncP-9 Naphthalene-Catabolic Plasmid NAH7 and Characterization of Its oriT Region and Relaxase. Appl. Environ. Microbiol. 83, (2017).

26. Fernández-López, C. et al. Functional properties and structural requirements of the plasmid pMV158-encoded MobM relaxase domain. J. Bacteriol. 195, 3000–3008 (2013).

27. Chen, Y., Staddon, J. H. & Dunny, G. M. Specificity determinants of conjugative DNA processing in the Enterococcus faecalis plasmid pCF10 and the Lactococcus lactis plasmid pRS01. Mol. Microbiol. 63, 1549–1564 (2007).

28. Jandle, S. & Meyer, R. Stringent and Relaxed Recognition of oriT by Related Systems for Plasmid Mobilization: Implications for Horizontal Gene Transfer. Journal of Bacteriology vol. 188 499–506 (2006).

29. Fernández-González, E. et al. A Functional oriT in the Ptw Plasmid of Burkholderia cenocepacia Can Be Recognized by the R388 Relaxase TrwC. Front Mol Biosci 3, 16 (2016).

30. Parker, C., Becker, E., Zhang, X., Jandle, S. & Meyer, R. Elements in the co-evolution of relaxases and their origins of transfer. Plasmid 53, 113–118 (2005).

31. Becker, E. C. & Meyer, R. J. Relaxed specificity of the R1162 nickase: a model for evolution of a system for conjugative mobilization of plasmids. J. Bacteriol. 185, 3538–3546 (2003).

32. O’Brien, F. G. et al. Origin-of-transfer sequences facilitate mobilisation of non-conjugative antimicrobial-resistance plasmids in Staphylococcus aureus. Nucleic Acids Res. 43, 7971–7983 (2015).

33. Ramsay, J. P. & Firth, N. Diverse mobilization strategies facilitate transfer of non-conjugative mobile genetic elements. Curr. Opin. Microbiol. 38, 1–9 (2017).

34. Pollet, R. M. et al. Processing of Nonconjugative Resistance Plasmids by Conjugation Nicking Enzyme of Staphylococci. J. Bacteriol. 198, 888–897 (2016).

35. Moran, R. A. & Hall, R. M. pBuzz: A cryptic rolling-circle plasmid from a commensal Escherichia coli has two inversely oriented oriTs and is mobilised by a B/O plasmid. Plasmid 101, 10–19 (2019).

36. Li, X. et al. oriTfinder: a web-based tool for the identification of origin of transfers in DNA sequences of bacterial mobile genetic elements. Nucleic Acids Res. 46, W229–W234 (2018).

37. Altschul, S. F., Gish, W., Miller, W., Myers, E. W. & Lipman, D. J. Basic local alignment search tool. J. Mol. Biol. 215, 403–410 (1990).

38. Williams, S. L. & Schildbach, J. F. TraY and integration host factor oriT binding sites and F conjugal transfer: sequence variations, but not altered spacing, are tolerated. J. Bacteriol. 189, 3813–3823 (2007).

39. Sut, M. V., Mihajlovic, S., Lang, S., Gruber, C. J. & Zechner, E. L. Protein and DNA effectors control the TraI conjugative helicase of plasmid R1. J. Bacteriol. 191, 6888–6899 (2009).

40. Frost, L. S., Ippen-Ihler, K. & Skurray, R. A. Analysis of the sequence and gene products of the transfer region of the F sex factor. Microbiol. Rev. 58, 162–210 (1994).

41. Kolomeisky, A. B. Physics of protein--DNA interactions: mechanisms of facilitated target search. Phys. Chem. Chem. Phys. 13, 2088–2095 (2011).

42. Rohs, R. et al. Origins of specificity in protein-DNA recognition. Annu. Rev. Biochem. 79, 233–269 (2010).

43. Rohs, R. et al. The role of DNA shape in protein–DNA recognition. Nature 461, 1248–1253 (2009).

44. Samee, M. A. H., Bruneau, B. G. & Pollard, K. S. A De Novo Shape Motif Discovery Algorithm Reveals Preferences of Transcription Factors for DNA Shape Beyond Sequence Motifs. Cell Systems vol. 8 27–42.e6 (2019).

45. Bansal, M., Kumar, A. & Yella, V. R. Role of DNA sequence based structural features of promoters in transcription initiation and gene expression. Curr. Opin. Struct. Biol. 25, 77–85 (2014).

46. Abeel, T., Saeys, Y., Bonnet, E., Rouze, P. & Van de Peer, Y. Generic eukaryotic core promoter prediction using structural features of DNA. Genome Research vol. 18 310–323 (2008).

47. Dao, F.-Y., Lv, H., Wang, F. & Ding, H. Recent Advances on the Machine Learning Methods in Identifying DNA Replication Origins in Eukaryotic Genomics. Front. Genet. 9, 613 (2018).

48. Chen, W., Feng, P. & Lin, H. Prediction of replication origins by calculating DNA structural properties. FEBS Lett. 586, 934–938 (2012).

49. Francia, M. V. et al. A classification scheme for mobilization regions of bacterial plasmids. FEMS Microbiol. Rev. 28, 79–100 (2004).

50. Shintani, M., Sanchez, Z. K. & Kimbara, K. Genomics of microbial plasmids: classification and identification based on replication and transfer systems and host taxonomy. Front. Microbiol. 6, 242 (2015).

51. Smillie, C. & Garcillan-Barcia, P. M., Victoria Francia, M., Rocha, EPC, and de la Cruz, F. (2010). Mobility of Plasmids. Microbiol. Mol. Biol. Rev. 74, 00020–00010.

52. Barabási, A.-L. & Oltvai, Z. N. Network biology: understanding the cell’s functional organization. Nat. Rev. Genet. 5, 101–113 (2004).

53. Shintani, M. et al. Single-cell analyses revealed transfer ranges of IncP-1, IncP-7, and IncP-9 plasmids in a soil bacterial community. Appl. Environ. Microbiol. 80, 138–145 (2014).

54. Klümper, U. et al. Broad host range plasmids can invade an unexpectedly diverse fraction of a soil bacterial community. ISME J. 9, 934–945 (2015).

55. Pignatelli, M., Moya, A. & Tamames, J. EnvDB, a database for describing the environmental distribution of prokaryotic taxa. Environ. Microbiol. Rep. 1, 191–197 (2009).

56. Lloyd-Price, J. et al. Strains, functions and dynamics in the expanded Human Microbiome Project. Nature 550, 61–66 (2017).

57. Human Microbiome Project Consortium. Structure, function and diversity of the healthy human microbiome. Nature 486, 207–214 (2012).

58. Escapa, I. F. et al. New Insights into Human Nostril Microbiome from the Expanded Human Oral Microbiome Database (eHOMD): a Resource for the Microbiome of the Human Aerodigestive Tract. mSystems 3, (2018).

59. Forster, S. C. et al. HPMCD: the database of human microbial communities from metagenomic datasets and microbial reference genomes. Nucleic Acids Res. 44, D604–9 (2016).

60. Dewhirst, F. E. et al. The human oral microbiome. J. Bacteriol. 192, 5002–5017 (2010).

61. Founou, L. L., Founou, R. C. & Essack, S. Y. Antibiotic Resistance in the Food Chain: A Developing Country-Perspective. Frontiers in Microbiology vol. 7 (2016).

62. Marshall, B. M., Ochieng, D. J. & Levy, S. B. Commensals: underappreciated reservoir of antibiotic resistance. Microbe Wash. DC 4, 231–238 (2009).

63. Hutchings, M. I., Truman, A. W. & Wilkinson, B. Antibiotics: past, present and future. Curr. Opin. Microbiol. 51, 72–80 (2019).

64. Manaia, C. M. Assessing the Risk of Antibiotic Resistance Transmission from the Environment to Humans: Non-Direct Proportionality between Abundance and Risk. Trends Microbiol. 25, 173–181 (2017).

65. Lanka, E. & Wilkins, B. M. DNA processing reactions in bacterial conjugation. Annu. Rev. Biochem. 64, 141–169 (1995).

66. Williams, S. L. & Schildbach, J. F. Examination of an inverted repeat within the F factor origin of transfer: context dependence of F TraI relaxase DNA specificity. Nucleic Acids Res. 34, 426–435 (2006).

67. Carballeira, J. D., González-Pérez, B., Moncalián, G. & de la Cruz, F. A high security double lock and key mechanism in HUH relaxases controls oriT-processing for plasmid conjugation. Nucleic Acids Res. 42, 10632–10643 (2014).

68. Vedantam, G., Knopf, S. & Hecht, D. W. Bacteroides fragilis mobilizable transposon Tn5520 requires a 71 base pair origin of transfer sequence and a single mobilization protein for relaxosome formation during conjugation. Mol. Microbiol. 59, 288–300 (2006).

69. de la Cruz, F., Frost, L. S., Meyer, R. J. & Zechner, E. L. Conjugative DNA metabolism in Gram-negative bacteria. FEMS Microbiol. Rev. 34, 18–40 (2010).

70. Gillings, M. R. Evolutionary consequences of antibiotic use for the resistome, mobilome and microbial pangenome. Front. Microbiol. 4, 4 (2013).

71. Chandler, M. et al. Breaking and joining single-stranded DNA: the HUH endonuclease superfamily. Nat. Rev. Microbiol. 11, 525–538 (2013).

72. Ramsay, J. P. et al. An updated view of plasmid conjugation and mobilization in Staphylococcus. Mob. Genet. Elements 6, e1208317 (2016).

73. Bukowski, M. et al. Prevalence of Antibiotic and Heavy Metal Resistance Determinants and Virulence-Related Genetic Elements in Plasmids of Staphylococcus aureus. Front. Microbiol. 10, 805 (2019).

74. Soucy, S. M., Huang, J. & Gogarten, J. P. Horizontal gene transfer: building the web of life. Nat. Rev. Genet. 16, 472–482 (2015).

75. Bengtsson-Palme, J., Kristiansson, E. & Larsson, D. G. J. Environmental factors influencing the development and spread of antibiotic resistance. FEMS Microbiol. Rev. 42, (2018).

76. Han, Z. et al. Signal transduction network motifs and biological memory. J. Theor. Biol. 246, 755–761 (2007).

77. Siefert, J. L. Defining the mobilome. Methods Mol. Biol. 532, 13–27 (2009).

78. Seyed-Allaei, H., Bianconi, G. & Marsili, M. Scale-free networks with an exponent less than two. Phys. Rev. E Stat. Nonlin. Soft Matter Phys. 73, 046113 (2006).

79. Perry, J. A. & Wright, G. D. The antibiotic resistance ‘mobilome’: searching for the link between environment and clinic. Front. Microbiol. 4, (2013).

80. Anderson, M. J. A new method for non-parametric multivariate analysis of variance. Austral Ecol. 26, 32–46 (2001).

81. Ye, C., Ji, G., Li, L. & Liang, C. detectIR: a novel program for detecting perfect and imperfect inverted repeats using complex numbers and vector calculation. PLoS One 9, e113349 (2014).

82. Zankari, E. et al. Identification of acquired antimicrobial resistance genes. J. Antimicrob. Chemother. 67, 2640–2644 (2012).

