## Supplementary Information for "Multiple plasmid origin-of-transfer substrates enable the spread of natural antimicrobial resistance to human pathogens"

### Table of contents

| | | |
|---|---|---|
| Supplementary Figures | … | p.2 - p.29 |
| Supplementary Tables | … | p.30 - p.46 |
| Supplementary References | … | p.47 - p.49 |

### Supplementary figures

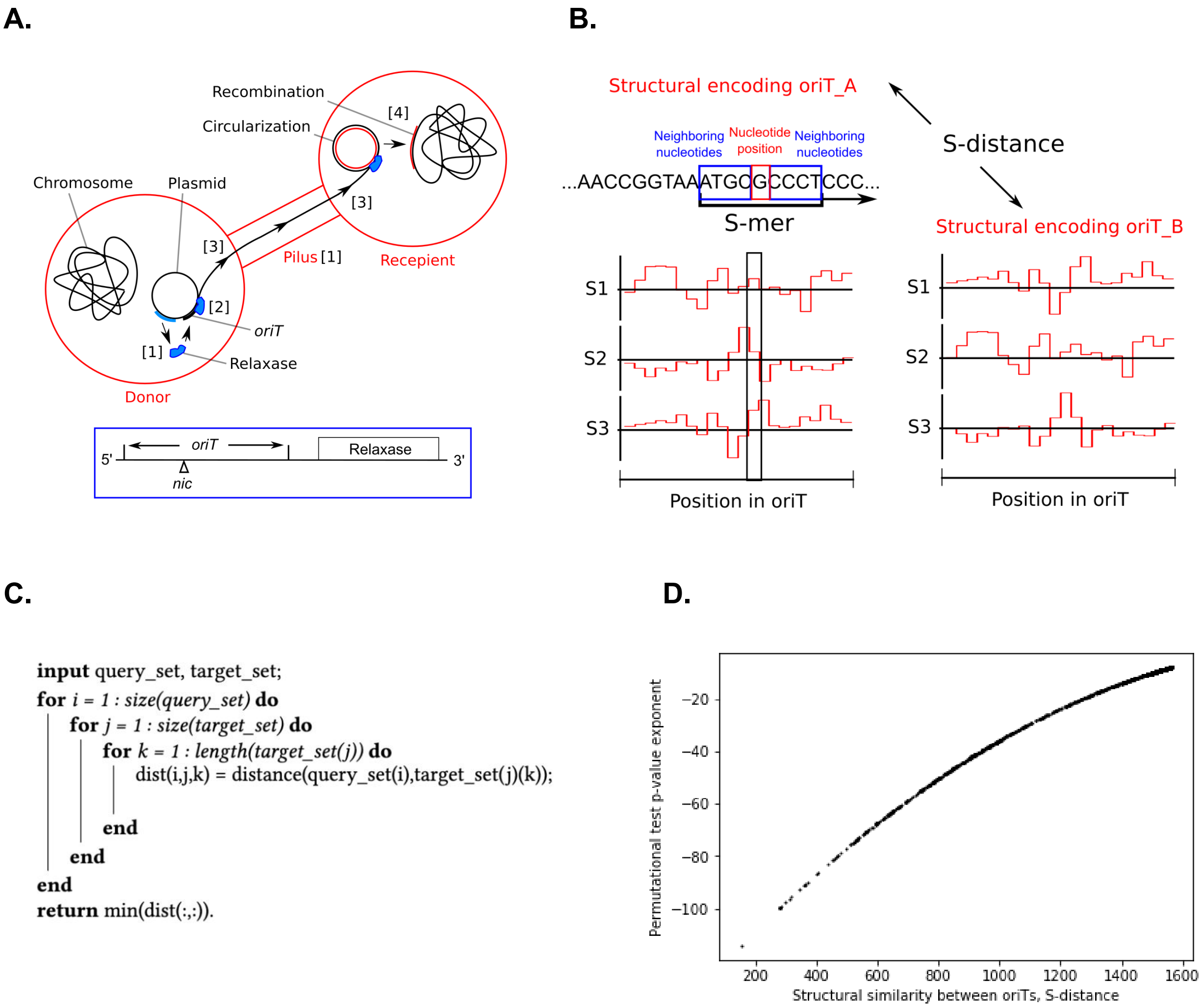


**Figure S1-1.** Overview of the structural alignment framework. (A) Depiction of the plasmid conjugation process, which can be divided into 4 steps: (i) formation of a conjugative pilus that connects the donor and recipient cells for transmission of mobile DNA, (ii) expression of enzymes (e.g. relaxase) and structural proteins, which recognize the binding sites at the DNA origin of transfer (oriT), where plasmid transfer is initiated, (iii) relaxase cuts into the oriT at the *nic* site, exposes the single-stranded DNA and, with the help of the protein transport system, transfers DNA to the recipient cell, (iv) in the recipient, either the missing DNA strand is synthesized and then circularized, in case of plasmid transfer, or the mobile DNA is integrated into the chromosome by recombinant mechanisms, whereas in the donor cell reconstruction of the missing DNA occurs. (B) Depiction of the DNA structural encoding and s-distance, where (i) consecutive k-mers of the DNA (k = 7) are encoded with clustered DNA structural property embeddings (marked s1, s2, s3; 18 such embeddings used) into a compressed representation termed 's-mers' (ii) the s-distance is the

Euclidean distance between all respective embeddings of two such structurally encoded DNA sequences. (C) Pseudocode giving an outline of the simplified alignment framework, which allows the use of different types of DNA distance measures between the target and query sequences, namely the structurally encoded s-distance and nucleotide sequence p-distance. (D) Mapping of s-distance scores to p-values obtained using permutational (bootstrap) tests, where bootstraps of the query oriT sequences were used to estimate p-values at cutoffs from 1e-6 to 1e-1. These points together with the theoretically predicted limit ~1e-132 were then used to fit to a second order polynomial function ($f = p_0 \cdot x^2 + p_1 \cdot x + p_2$ ; $p_0 = -3.045e-05$ ,$p_1 = 0.128$, $p_2 = -133.000$).

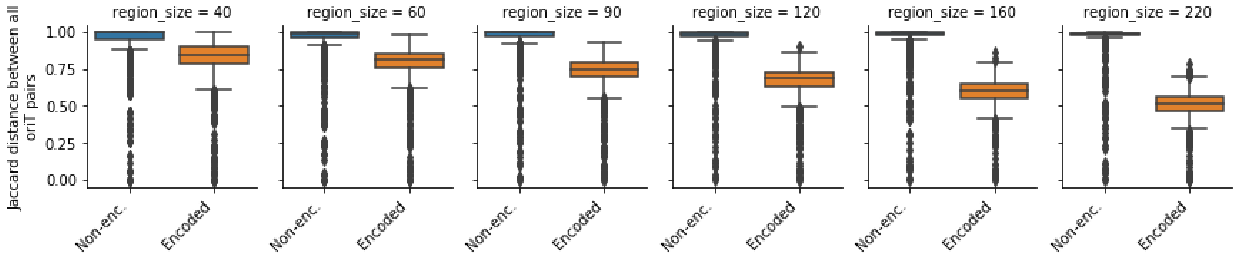


**Figure S1-2.** Distribution of pairwise Jaccard distances between oriTs, using structurally encoded k-mers (Methods M2) or non-encoded nucleotide k-mers, with the subsets of different oriT sizes.

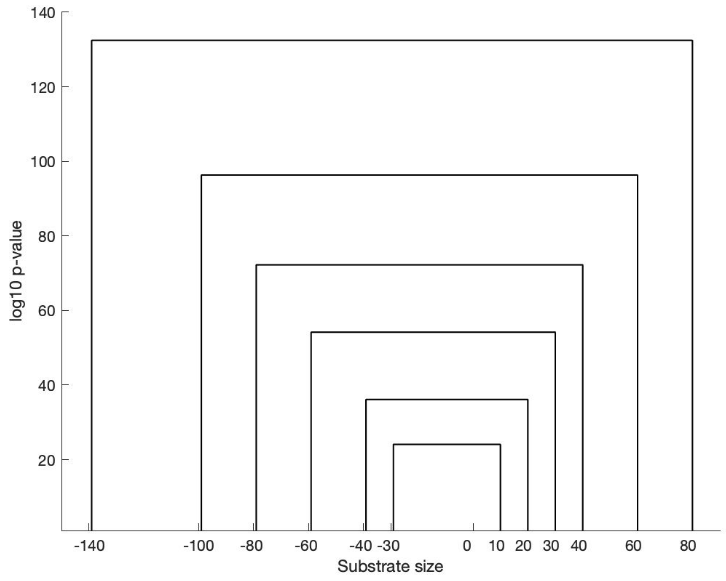


**Figure S1-3.** Schematic diagram of the estimated maximum statistical depth achievable with different sequence lengths, where an over 1e100-fold difference is observed between the 40 bp and 220 bp sizes of oriT.

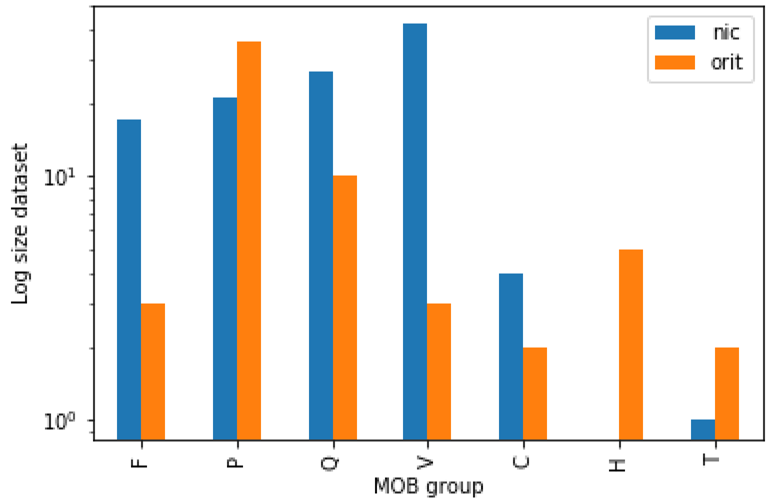


**Figure S1-4.** Size of plasmid datasets with known oriT regions across 7 MOB groups. 65% of elements were used as query sequences as locations of *nic* sites (meaning also oriT regions) were known, whereas the rest with experimentally or putatively determined oriT region locations but unknown nic sites were used for testing. For development and testing of the structural alignment algorithm only Mob groups {F,P,Q,V} had sufficient data, ie. over 10 nic sites known per Mob group, which resulted in datasets of 106 query regions (nic and mob known), 106 query plasmids used for 10-fold cross validation tests (nic and mob known)] and 51 testing plasmids (oriT and mob known). The full query dataset comprising 7 Mob groups contained 112 distinct oriT sequences.

**A.**

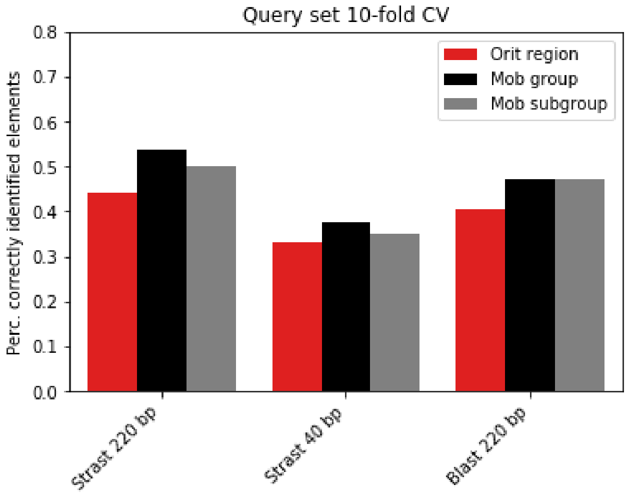


**B.**

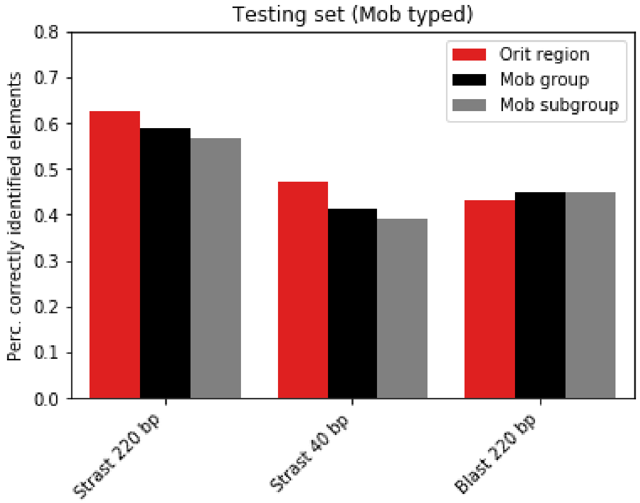


**Figure S1-5.** Percentage of correctly identified elements using our structural alignment algorithm (Strast) with 220 bp and 40 bp query regions sizes and Blast with 220 bp region sizes with (A) query dataset 10-fold cross-validations and (B) Mob-typed testing dataset.

**A.**

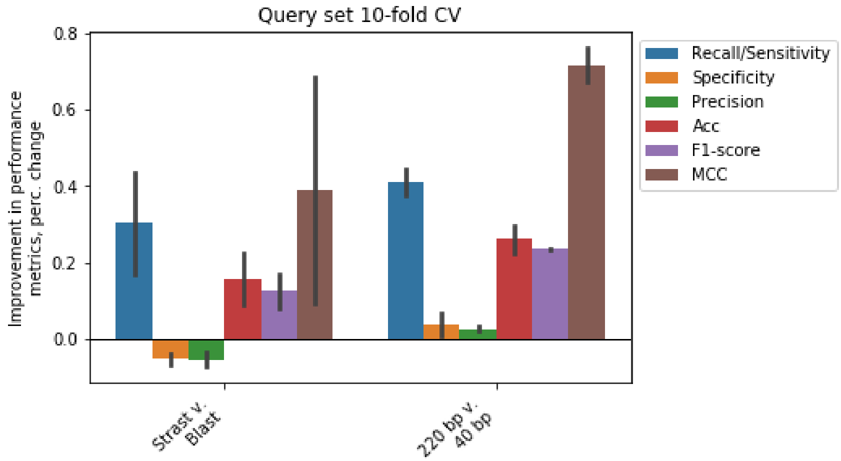


**B.**

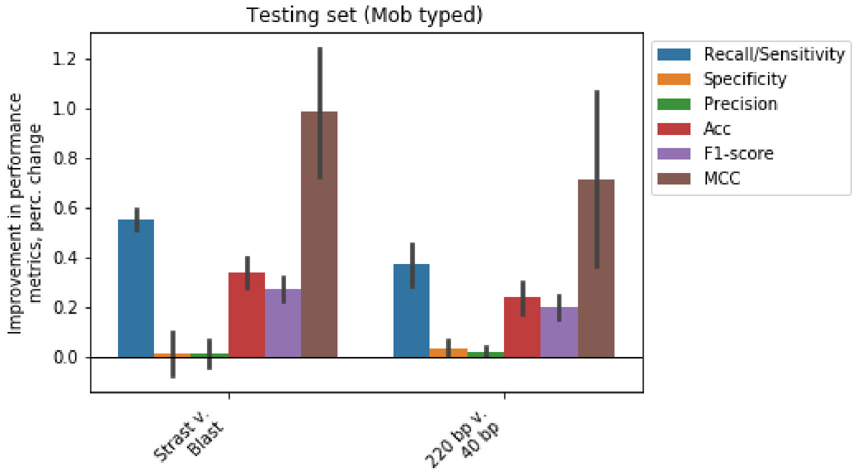


**Figure S1-6.** Relative changes in the six measured performance metrics averaged across oriT identification and Mob typing with (A) query dataset 10-fold cross-validations and (B) Mob-typed testing dataset. Error bars denote 95% confidence intervals.

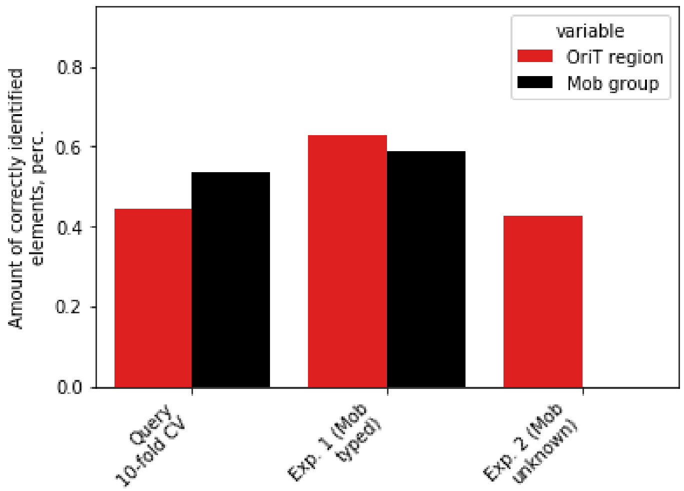


**Figure S1-7.** Amounts of correctly identified elements across the query and two testing datasets with known or unknown MOB groups, for oriT typing as well as discrimination of MOB groups and subgroups.

**A.**

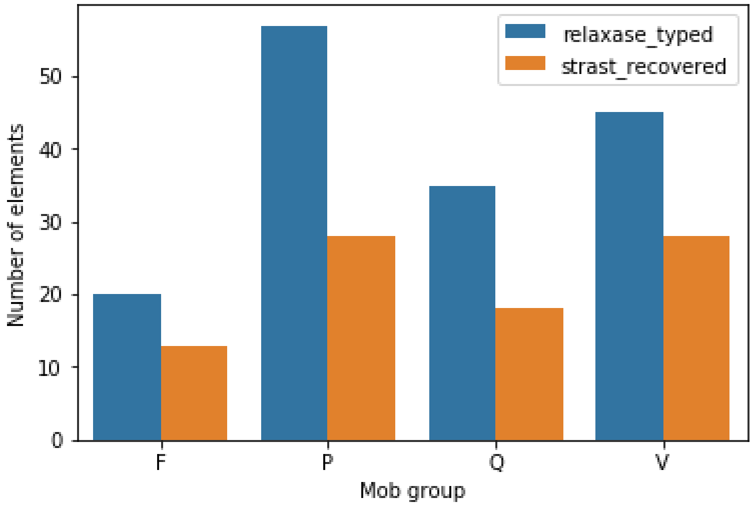


**B.**

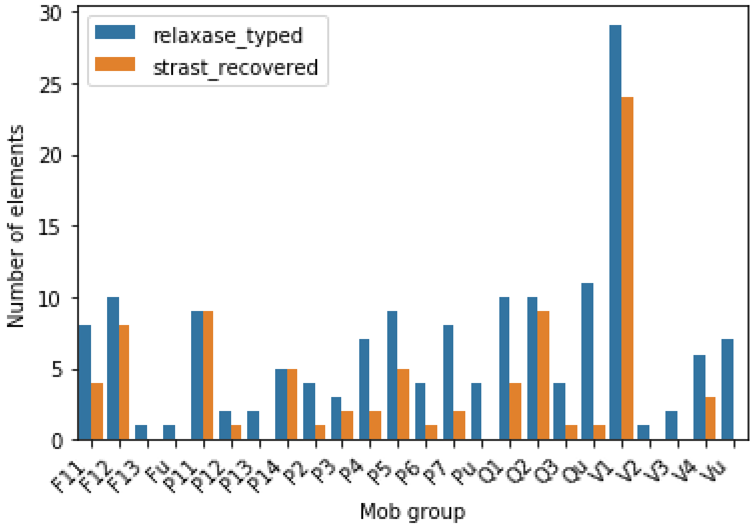


**Figure S1-8.** Number of recovered elements across (A) 4 Mob groups and (B) 24 Mob subgroups in the combined testing datasets (10-fold cross-validations on query plasmids and 51 Mob-typed plasmids).

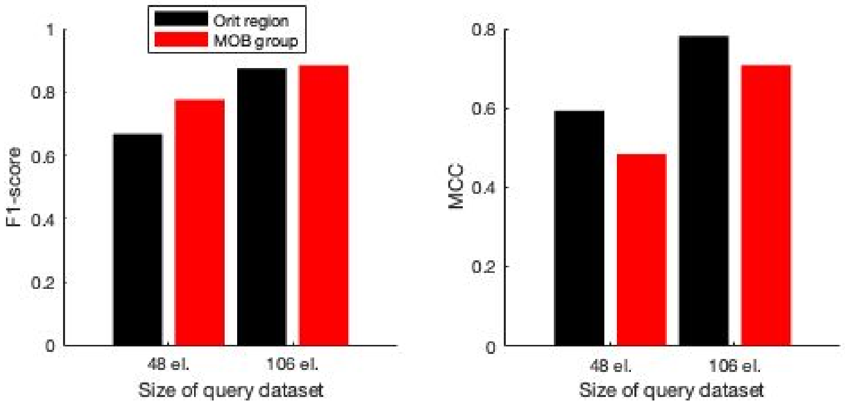


**Figure S1-9.** Performance metrics F1-score and Matthews correlation coefficient (MCC) showing the effect of query dataset size with a diluted set of 48 elements compared to the full query dataset.

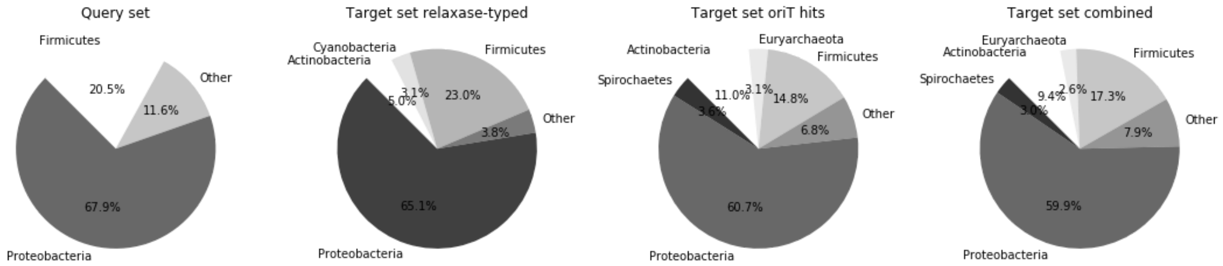


**Figure S2-1.** Distributions of phyla in the query dataset as well as in the target dataset obtained by relaxase and structural alignment-typing.

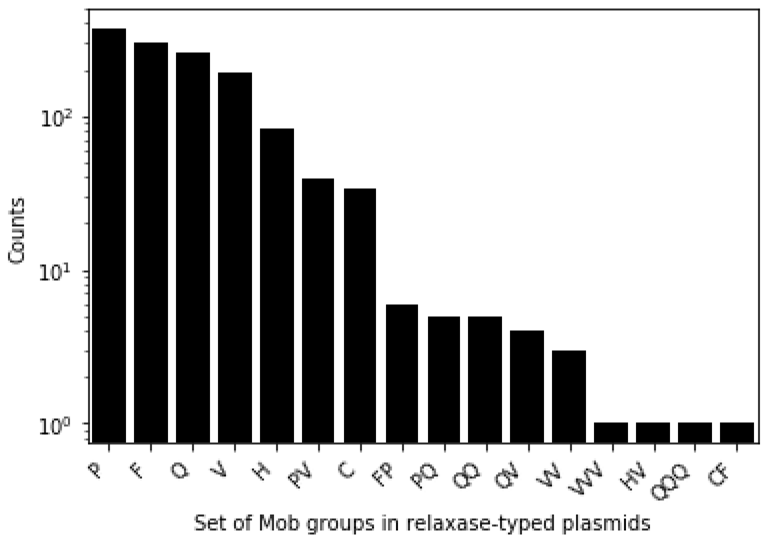


**Figure S2-2.** Distributions of single and sets of Mob groups in relaxase-typed plasmids in the target dataset.

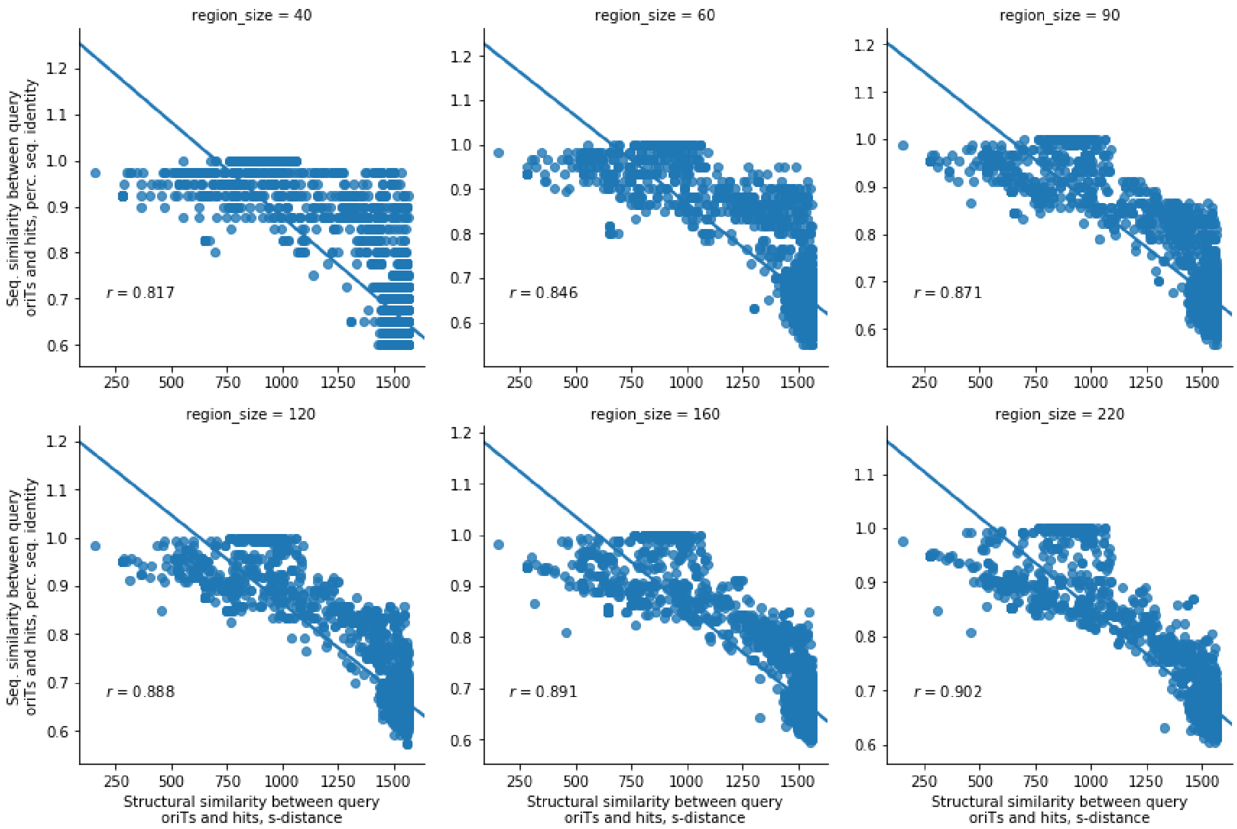


**Figure S2-3.** Correlation analysis between the sequence homology and structural similarities (s-distance) among oriT hits and their closest-associated query sequences. All p-values were below 1e-16.

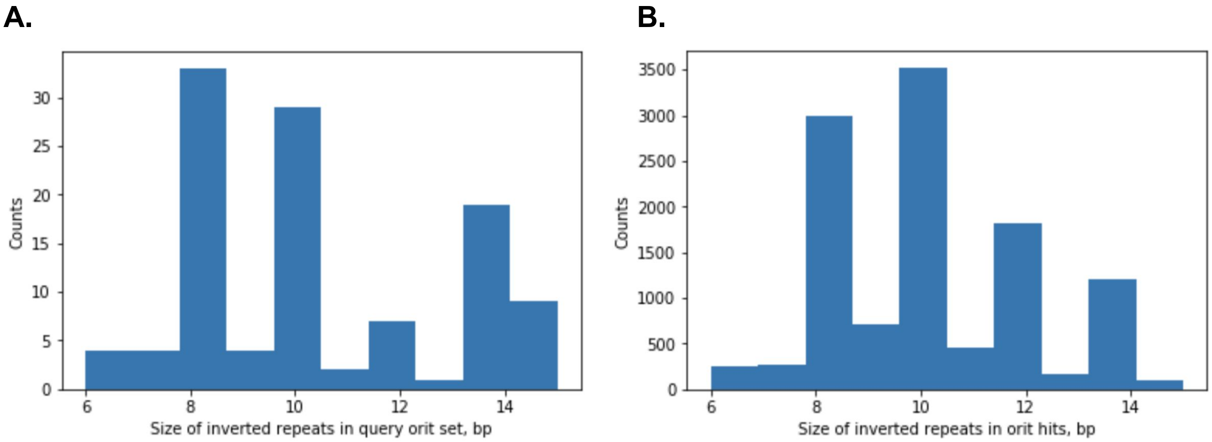


**Figure S2-4.** Distributions of sizes of inverted repeats identified in the (A) query dataset and (B) dataset of oriT hits.

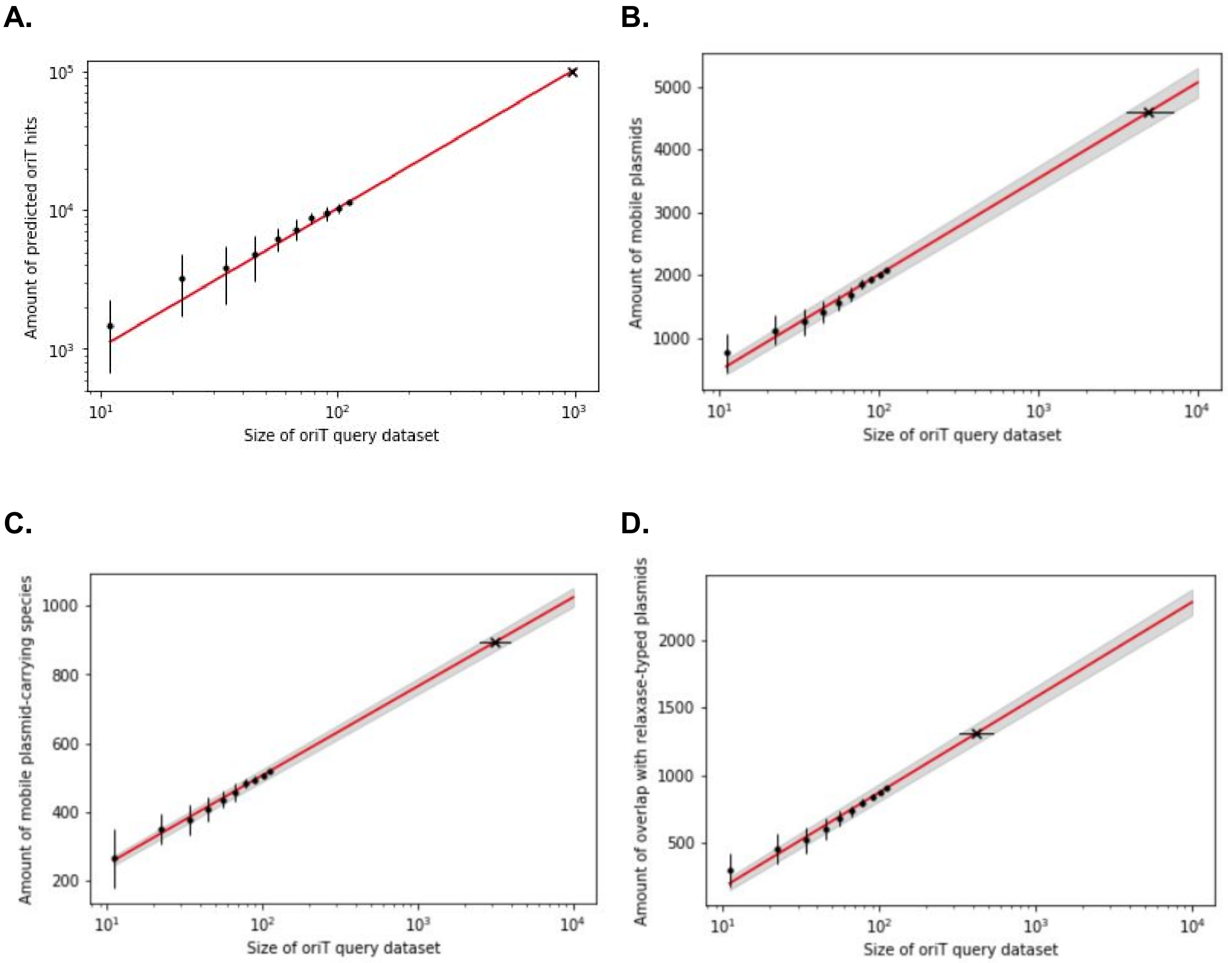


**Figure S2-5.** Simulated effect of the size of query dataset on the (A) amount of uncovered oriT regions, (B) amount of mobile plasmids, (C) amount of mobile plasmid-carrying host species and (D) amount of overlap with relaxase-typed plasmids. Red lines denote least squares fit and gray areas denote 95% confidence intervals. Black dots and error bars denote 10 repetitions of 10-fold dataset dilutions used for curve fitting. 'X' and vertical error bars denote predictions that mark the size of query dataset required to recover (A) 1e5 oriT hits (query dataset of 975 oriTs with 95% lower and upper bounds within 0.07 of this value, respectively), (B) OriT regions spanning the whole target dataset of 4602 plasmids (query dataset of 4940 oriTs, 95% lower and upper bounds were 3548 and 7003, respectively), (C) the whole species diversity of the target dataset - 893 unique species (query dataset of 3101 oriTs, 95% lower and upper bounds were 2491 and 3891), (D) a full overlap with the relaxase-typed plasmids (query dataset of 415 oriTs, 95% lower and upper bounds were 328 and 532, respectively). Horizontal and vertical error bars denote 95 % confidence intervals.

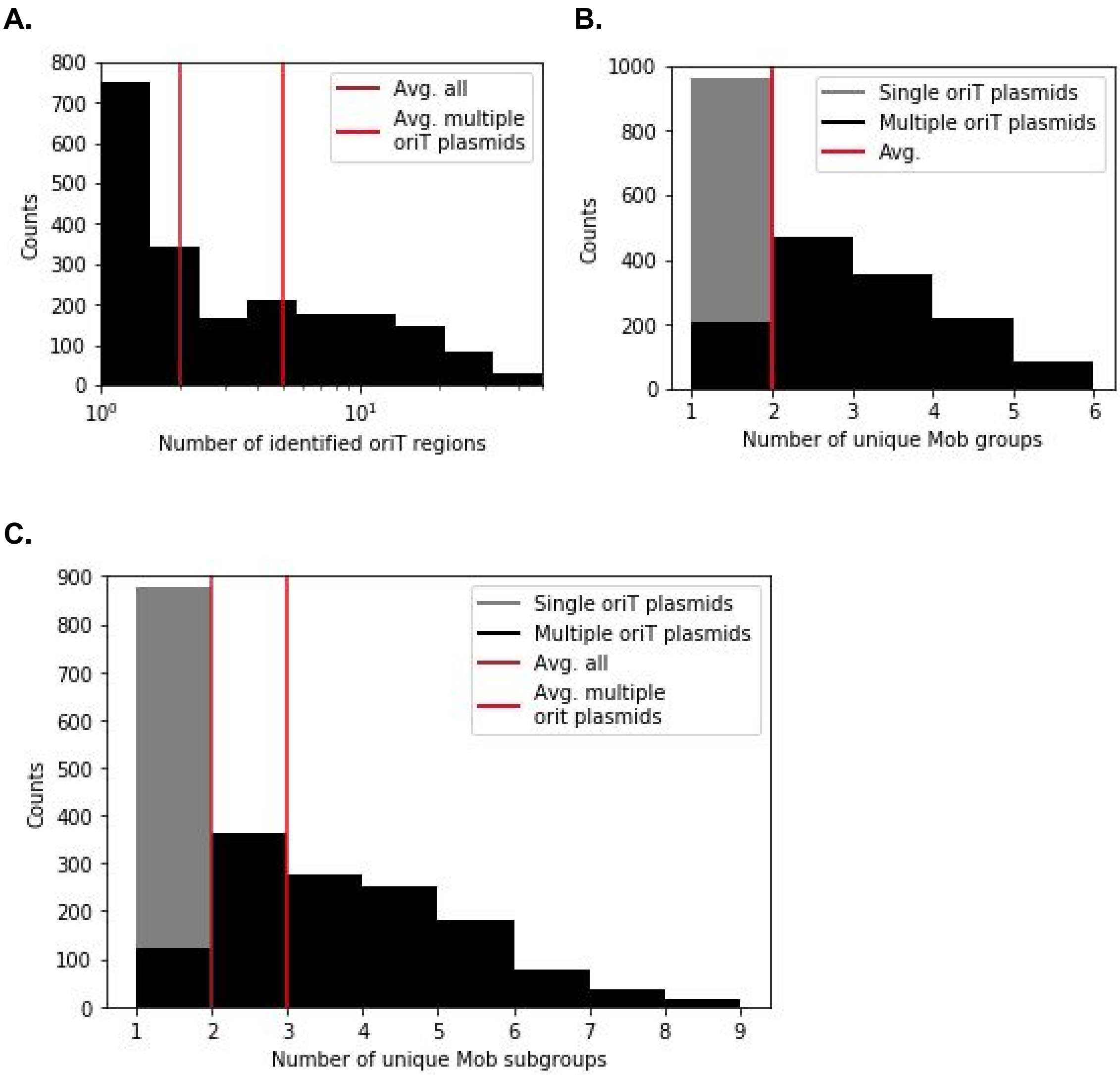


**Figure S3-1.** (A) Distribution of the amount of identified oriTs. Separate median averages are given for all mobile plasmids and those with multiple oriTs. (B) Distribution of the number of unique Mob groups in single and multiple oriT plasmids. Median averages for all mobile plasmids and those with multiple oriTs are the same. (C) Distribution of the number of unique Mob subgroups in single and multiple oriT plasmids. Separate median averages are given for all mobile plasmids and those with multiple oriTs.

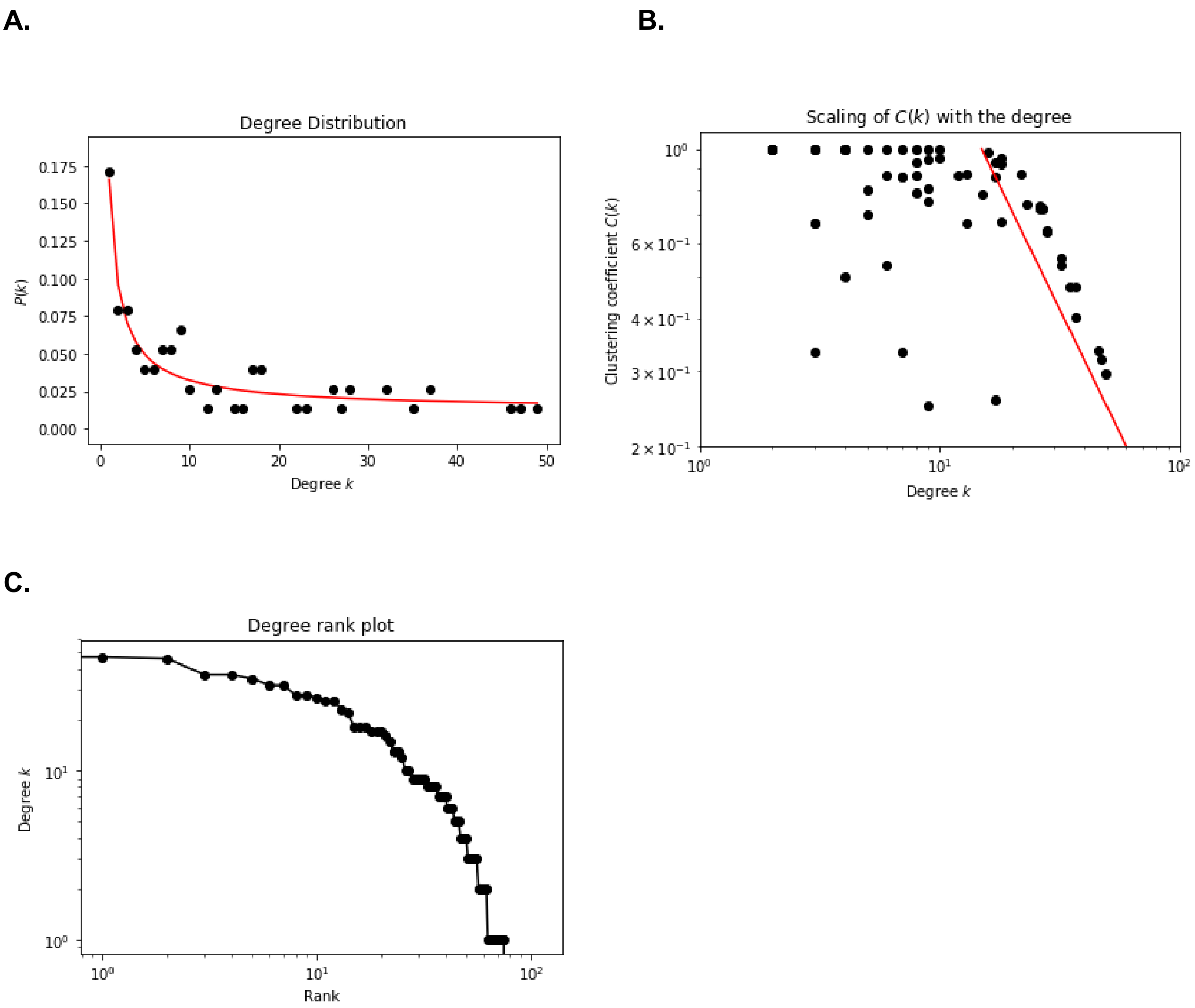


**Figure S3-2.** (A) Degree distribution with the power law function ($x^{-a} \cdot b + c$) fit to the data using least squares regression (red line). Exponents were 0.965, 0.147, 0.012, respectively. The characteristic of networks with degree exponents below 2 is that the average degree grows with their size (Seyed-Allaei, Bianconi, and Marsili 2006). (B) Scaling of clustering coefficient with the degree of connectivity per node. Red line denotes approximate linear fit. (C) Degree rank plot.

**A.**

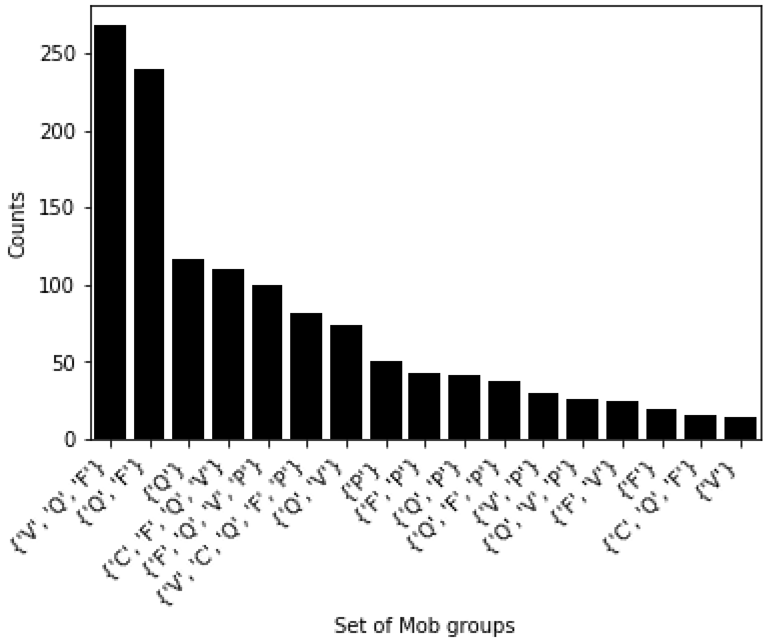


**B.**

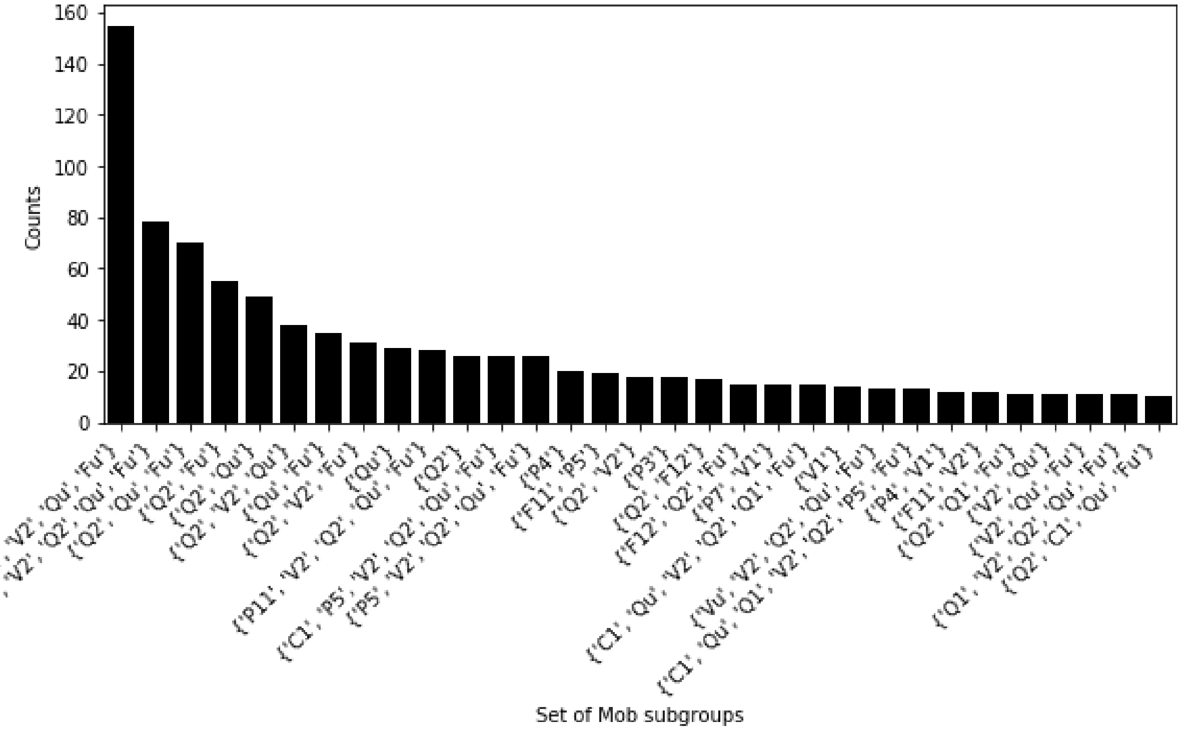


**Figure S3-3.** Most frequent sets of (A) Mob groups and (B) Mob subgroups with a number of occurrences 10 or above.

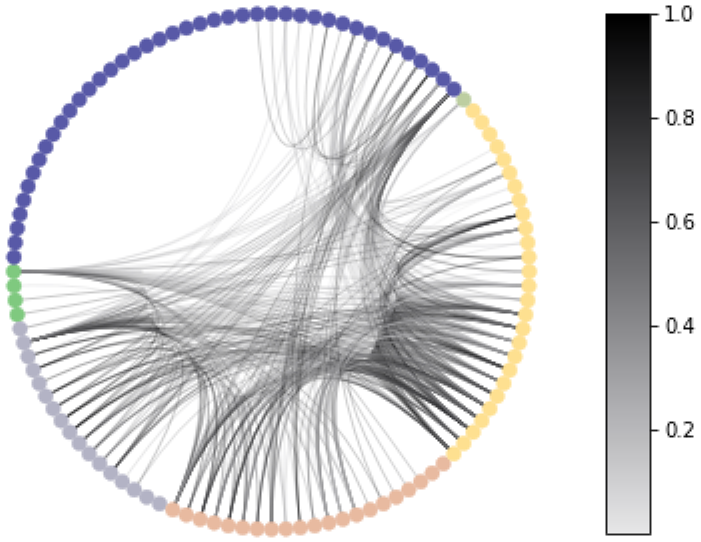


**Figure S3-4.** Circos plot of connections between Mob groups by parent (query) oriT regions found on the same plasmid. The color grey corresponds to Mob group F, orange to P, yellow to Q, blue to V, green to C and light gray to T. Strength of connections depicts the *p*-value from 1e-8 as 0 to 1e-99 as 1.

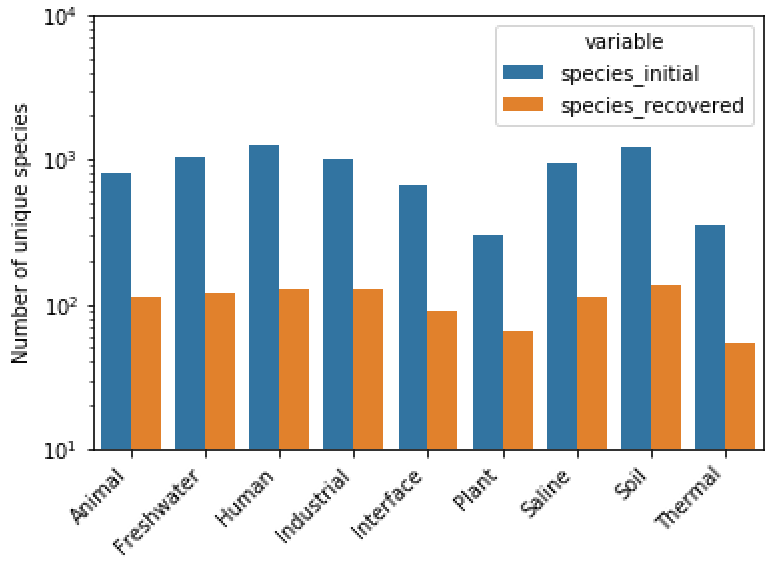


**Figure S4-1.** Number of unique species across habitats in the initial whole dataset and the recovered dataset obtained by merging plasmids and orit hits with the habitat data. A database of 16,072 mappings between microbial taxonomies and 9 habitat supertypes was obtained according to published data (Pignatelli, Moya, and Tamames 2009; Lloyd-Price et al. 2017; Human Microbiome Project Consortium 2012; Escapa et al. 2018; Forster et al. 2016; Dewhirst et al. 2010) (Tables S4-1 & S4-2). On average, merely 13% of the original species in the habitats was recovered by the mobile plasmid data (Fig S4-X), with an overall even distribution varying less than 3% around the average value (from 10.2% to 15.6%). Therefore the habitat sizes reflected those in the original habitat data (on average 939 species) but were almost 8-fold smaller (on average 119 species).

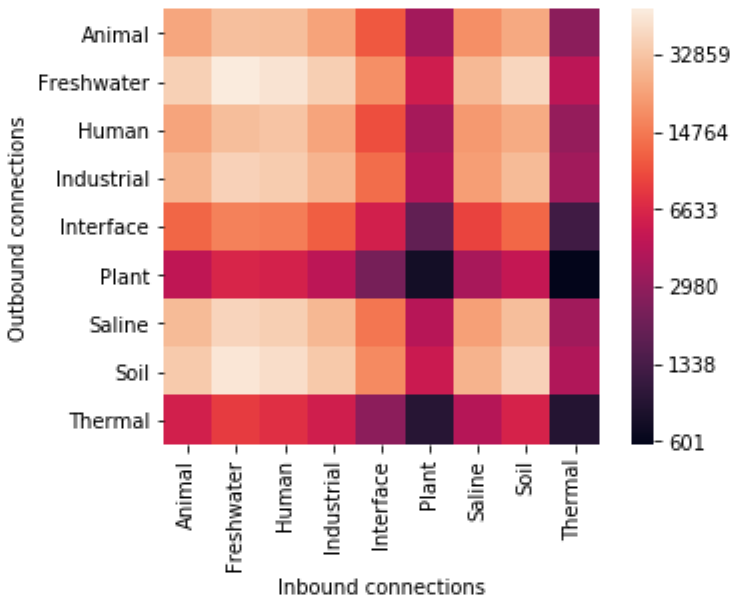


**Figure S4-2.** The connectivity of habitats (nodes) based on potential plasmid transfers (edges) from Fig 4A depicted as an adjacency matrix.

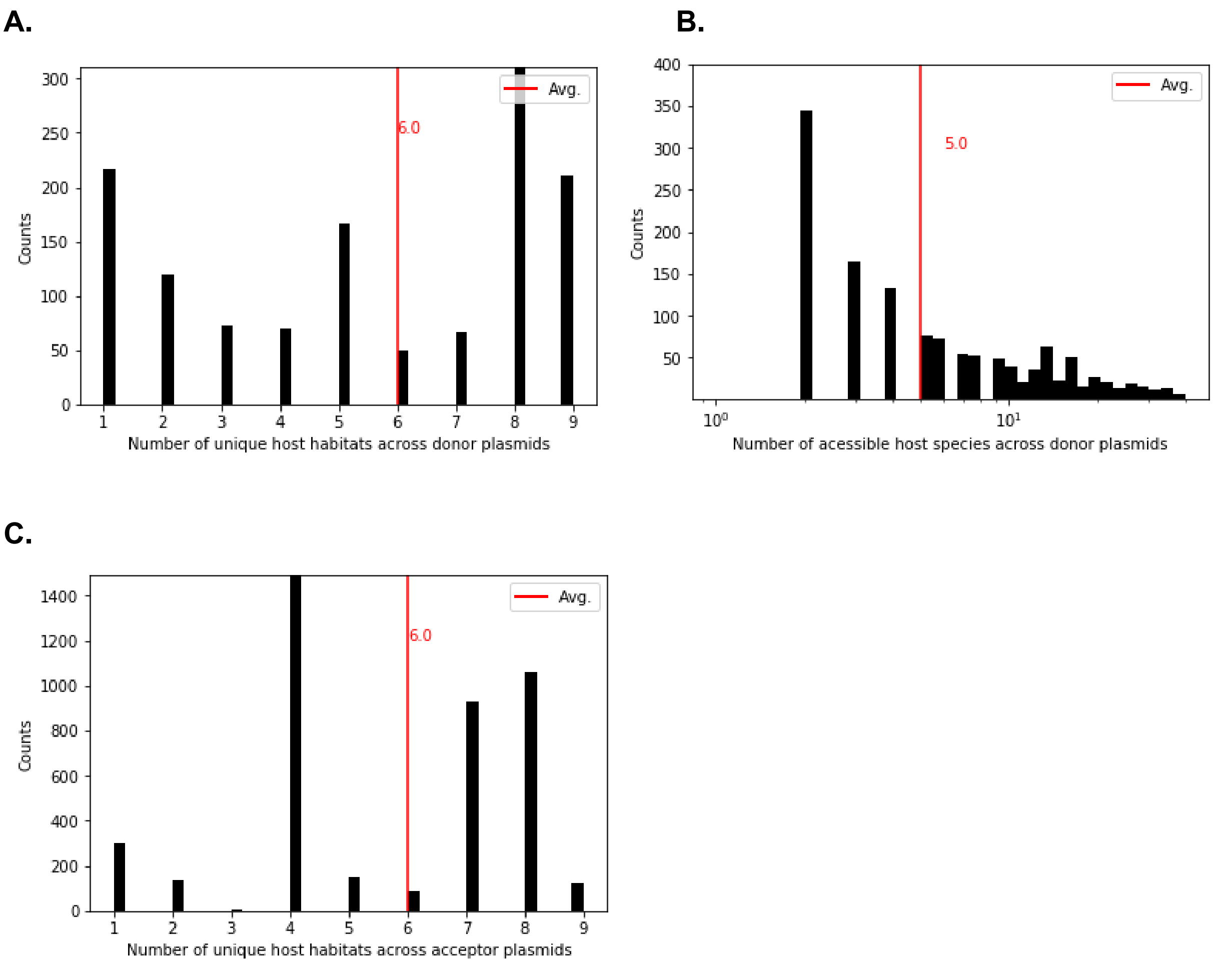


**Figure S4-3.** Properties of the horizontal transfer network. (A) Distribution of the number of unique host habitats per plasmid. (B) Distribution of the number of potentially accessible hosts of the multi-oriT donor plasmids. (C) Distribution of the number of unique accessible host habitats via oriTs.

**A.**

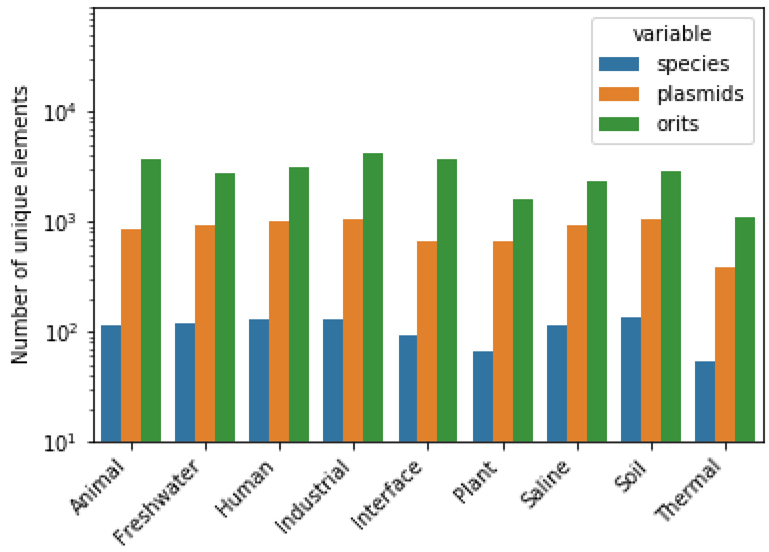


**B.**

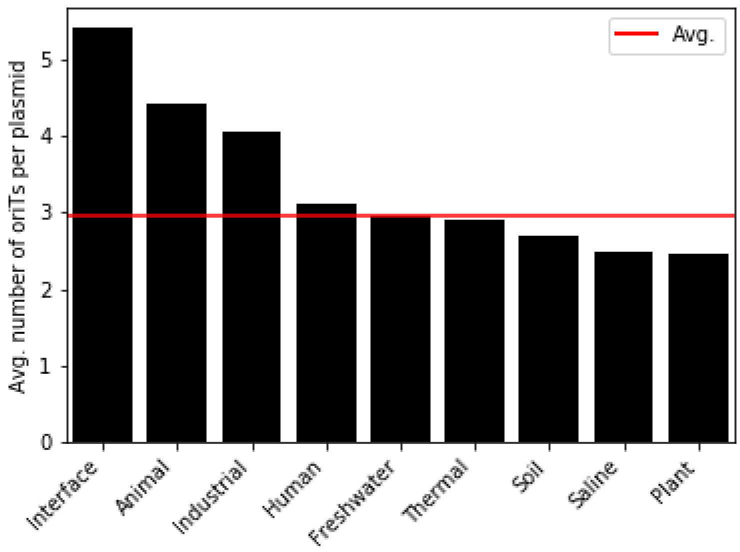


**Figure S4-4.** (A) Number of unique species, plasmids and oriT regions across habitats and (B) average number of oriTs per plasmid across habitats.

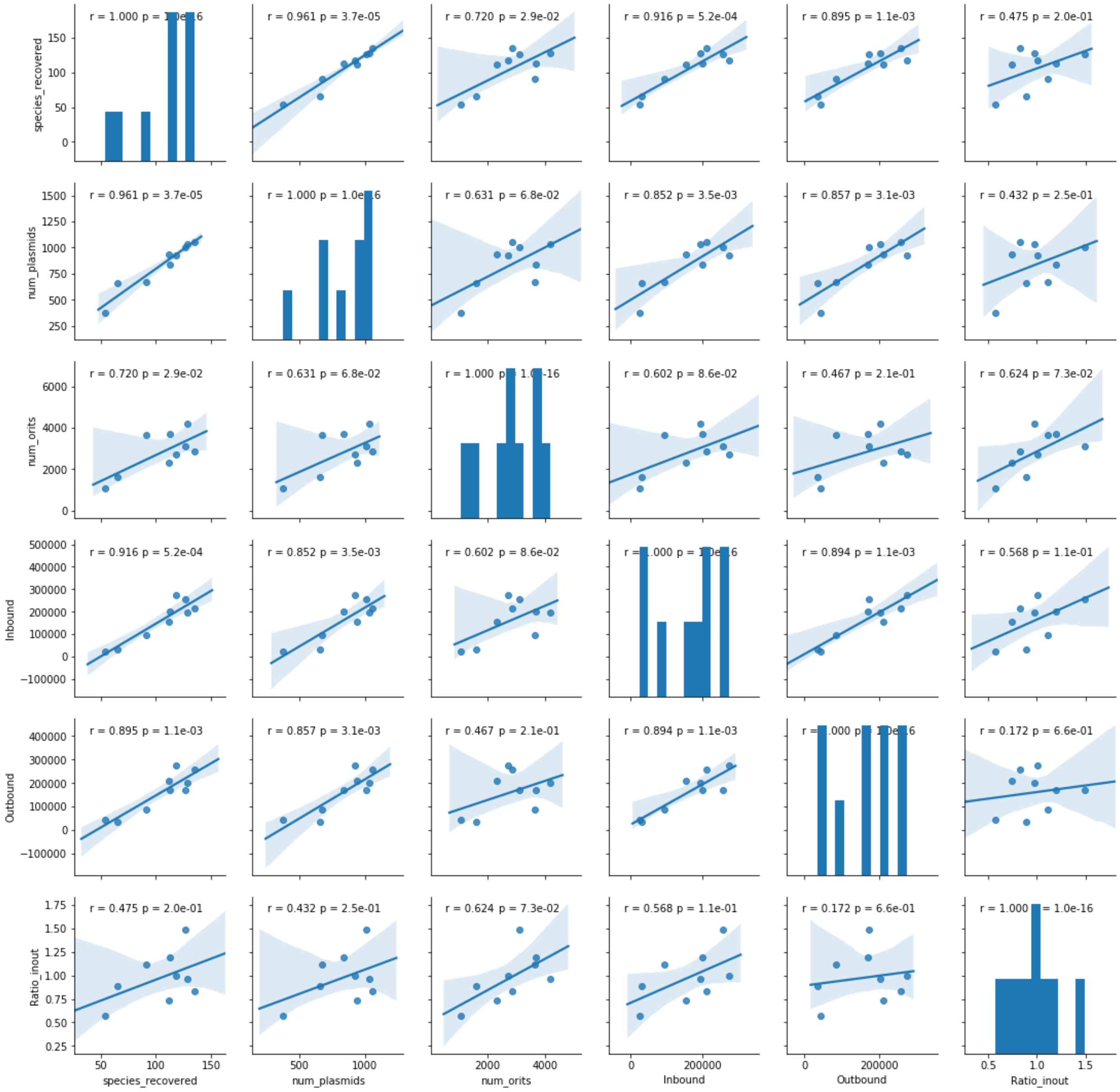


**Figure S4-5.** Correlation analysis between the number of unique species, plasmids and oriTs, inbound and outbound connections as well as the ratio between the number of inbound and outbound connections.

**A.**

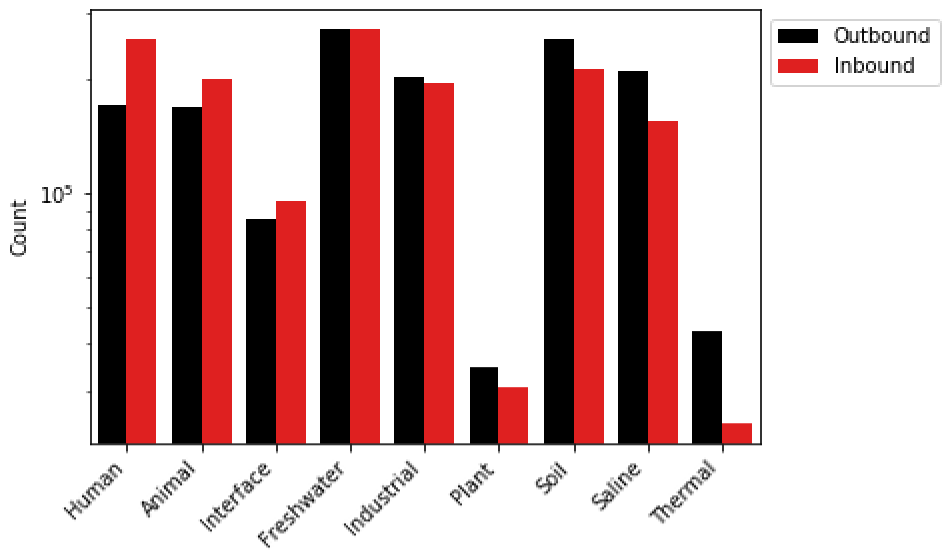


**B.**

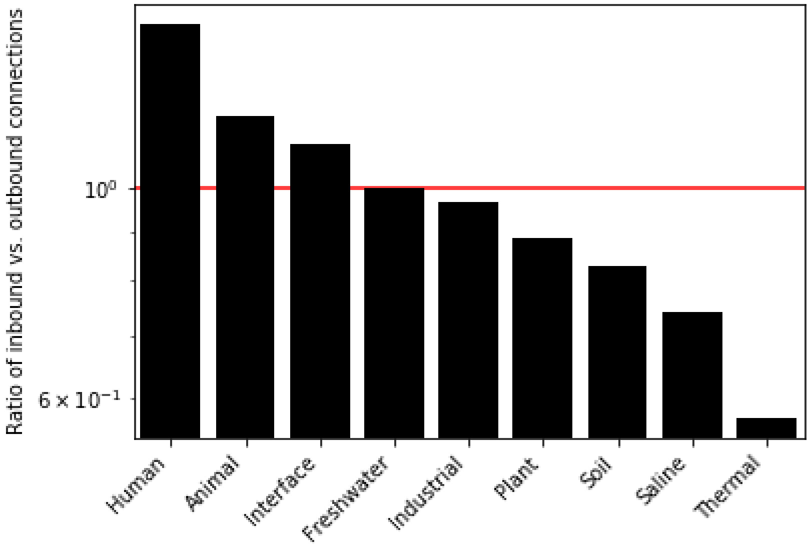


**Figure S4-6.** (A) Number of outbound and inbound connections and (B) ratio of inbound vs. outbound connections across the different habitats.

**A.** **B.**

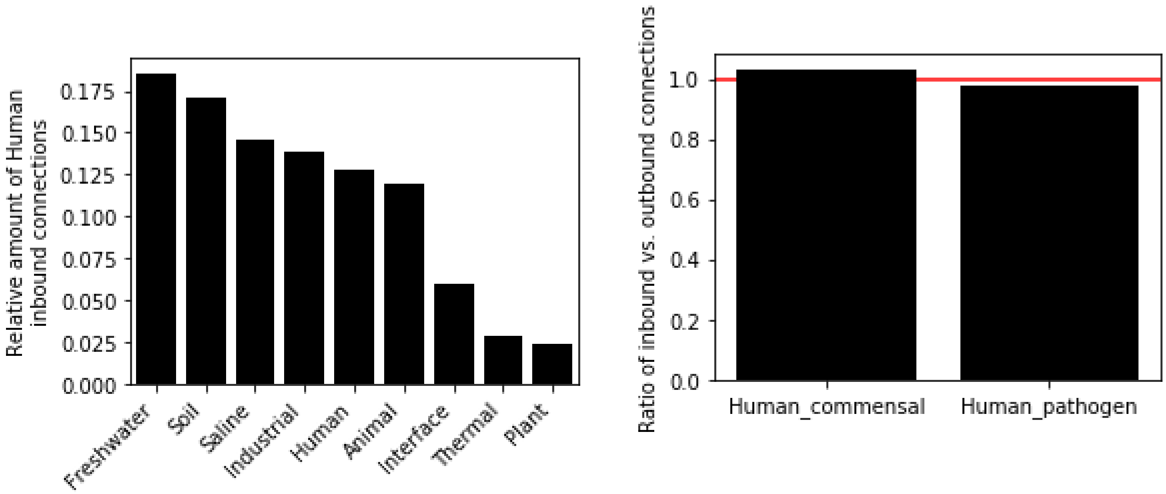


**Figure S4-7.** (A) Relative amount of connections to Human habitat from other environments. (B) Ratio of inbound vs. outbound connections observable within the human system.

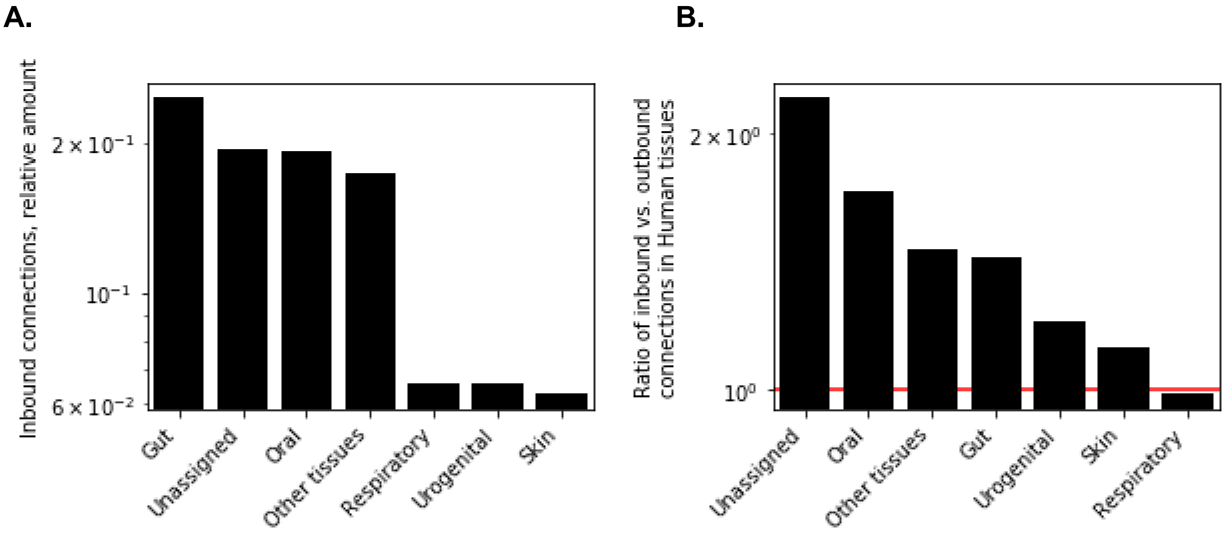


**Figure S4-8.** (A) Relative amount of inbound connections to the different human tissues. (B) Ratio of inbound vs. outbound connections across the human tissues.

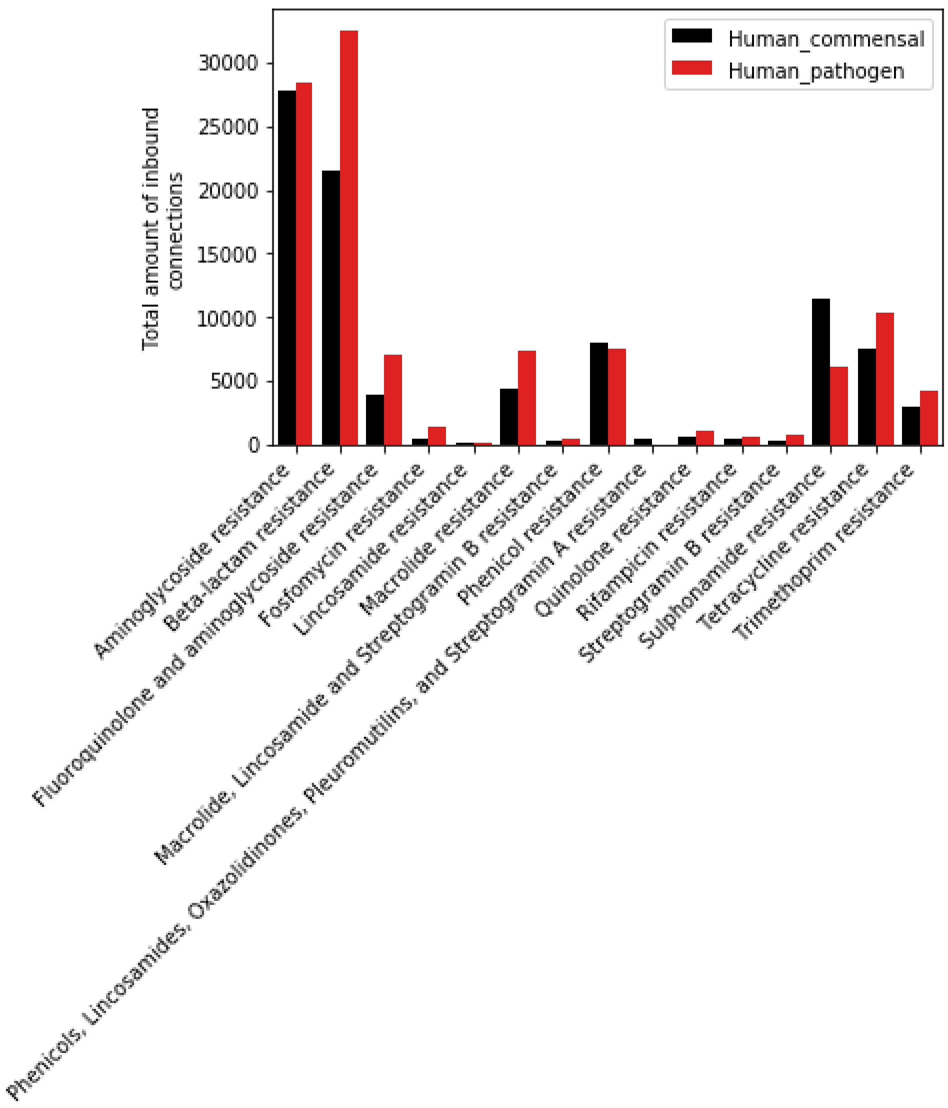


**Figure S4-9.** Amount of inbound connections of different AMR classes to the Human microbiome.

### Supplementary tables

**Table S1-1.** Datasets used for algorithm development.

| Development | Dataset | Groups | Availability/Reference |
|---|---|---|---|
| s-distance testing | Balanced dataset of 64 oriTs | 64 positive oriTs and 64 negative sequences | (Zrimec and Lapanje 2018) |
| s-distance testing | Balanced dataset of 64 oriTs | 4 Mob groups | (Zrimec and Lapanje 2018) |
| Alignment query | 106 oriTs with known nic sites | 4 Mob groups | Dataset S1 |
| Alignment testing 10-fold cross-validations | Plasmid sequences of the 106 query oriTs | 4 Mob groups | Dataset S1 |
| Alignment experimental testing 1 | 51 plasmids with known oriT regions, unknown nic sites | 4 Mob groups | Dataset S2 |
| Alignment experimental testing 2 | 13 plasmids with known nic sites | unknown Mob groups | (Li et al. 2018), Dataset S3 |

**Table S1-2.** Uncovered oriT regions in experimentally determined plasmids

| plasmid_name | query_orit | nic_location | mob | mob_sub group | p_value exponent | nic_oritdb | nic |
|---|---|---|---|---|---|---|---|
| pMAB01 | R751/pTP6 | 32985 | P | 11 | -36.6478 | CATCCTG\|C | TCCTGCCCGC |
| pSU233 | P307 | 138 | F | 12 | -52.3715 | GTGGGGTGT\|GG | GGGTGTGGTG |
| pMAS2027 | R6K | 38375 | P | 3 | -19.699 | TATCCTG\|C | CCTGCATCGC |
| pMAS2027 | R6K | 2066 | P | 3 | -13.3076 | TATCCTG\|C | ATCCTGCATC |
| pRJ6 | pC223 | 2185 | P | 7 | -12.9163 | TGCTTG\|CCA | TGCTTGCCAA |
| ICEKp1 | p29930 | 66528 | C | 1 | -12.1504 | GGTTG\|GTCGCG | GTTGGTCGCG |

**Table S1-3.** Models of physicochemical and conformational DNA properties.

| Variable name | Units | Model | Reference |
|---|---|---|---|
| dG | kcal/mol | thermodynamic | (SantaLucia 1998) |
| dH | kcal/mol | thermodynamic | (SantaLucia 1998) |
| dS | mol/K cal | thermodynamic | (SantaLucia 1998) |
| dGst | kcal/mol | thermodynamic | (Protozanova, Yakovchuk, and Frank-Kamenetskii 2004) |
| dGbp | kcal/mol | thermodynamic | (Protozanova, Yakovchuk, and Frank-Kamenetskii 2004) |
| dGkl | kcal/mol | thermodynamic | (Protozanova, Yakovchuk, and Frank-Kamenetskii 2004) |
| Tm | deg C | thermodynamic | (Gotoh and Tagashira 1981) |
| BC | kJ/mol | thermodynamic | (Breslauer et al. 1986) |
| BA | kJ/mol | thermodynamic | (Aida 1988) |
| BZ | kJ/mol | thermodynamic | (Ho et al. 1986; Hartmann, Malfoy, and Lavery 1989) |
| BA_k | kcal/mol | thermodynamic | (Kulkarni and Mukherjee 2013) |
| dGst_2 | kcal/mol | thermodynamic | (Perez et al. 2004) |
| Zp | A | unknown | (Kulkarni and Mukherjee 2013) |
| Twist | deg | curvature | (Perez et al. 2004) |
| Tilt | deg | curvature | (Perez et al. 2004) |
| Roll | deg | curvature | (Perez et al. 2004) |
| Shift | A | curvature | (Perez et al. 2004) |
| Slide | A | curvature | (Perez et al. 2004) |
| Rise | A | curvature | (Perez et al. 2004) |
| Phi_slide | kJ mol-1 A-2 | curvature | (Packer, Dauncey, and Hunter 2000) |
| Phi_shift | kJ mol-1 A-2 | curvature | (Packer, Dauncey, and Hunter 2000) |
| Maj_bend | mu | curvature | (Gartenberg and Crothers 1988) |
| Min_bend | mu | curvature | (Gartenberg and Crothers 1988) |
| Wdg | deg | wedge | (Bolshoy and McNamara 1991) |
| Dir | deg | wedge | (Bolshoy and McNamara 1991) |
| HT_wdg | deg | curvature | (Kabsch, Sander, and Trifonov 1982) |
| ProT2 | deg | curvature | (Gorin, Zhurkin, and Olson 1995) |
| C2 | A | clash function | (Gorin, Zhurkin, and Olson 1995) |
| MajS | A | clash function | (Gorin, Zhurkin, and Olson 1995) |
| MajD | A | clash function | (Gorin, Zhurkin, and Olson 1995) |
| MinS | A | clash function | (Gorin, Zhurkin, and Olson 1995) |

| | | | |
|---|---|---|---|
| MinD | A | clash function | (Gorin, Zhurkin, and Olson 1995) |
| z | nm | inverse | (Geggier and Vologodskii 2010) |
| h | bp/turn | NN | (Geggier and Vologodskii 2010) |
| z2_set1 | nm | inverse | (Sivolob and Khrapunov 1995) |
| z2_set2 | nm | inverse | (Sivolob and Khrapunov 1995) |
| Deform | deg^3 A^3 | NN | (Olson et al. 1998) |
| Twist2 | deg | curvature | (Olson et al. 1998) |
| Tilt2 | deg | curvature | (Olson et al. 1998) |
| Roll2 | deg | curvature | (Olson et al. 1998) |
| Shift2 | A | curvature | (Olson et al. 1998) |
| Slide2 | A | curvature | (Olson et al. 1998) |
| Rise2 | A | curvature | (Olson et al. 1998) |
| u2 | cleavage freq. | DNAzeI | (Ivan Brukner et al. 1995; I. Brukner et al. 1995) |
| Twist3 | deg | curvature | (Karas et al. 1996) |
| Rise3 | A | curvature | (Karas et al. 1996) |
| Bend | deg | curvature | (Karas et al. 1996) |
| Tip | deg | curvature | (Karas et al. 1996) |
| Inclination | deg | curvature | (Karas et al. 1996) |
| MajWidth | A | curvature | (Karas et al. 1996) |
| MajDepth | A | curvature | (Karas et al. 1996) |
| MinWidth | A | curvature | (Karas et al. 1996) |
| MinDepth | A | curvature | (Karas et al. 1996) |
| u | cleavage freq. | DNAzeI | (Ivan Brukner et al. 1995; I. Brukner et al. 1995) |
| var | fraction | nucleosome | (Satchwell, Drew, and Travers 1986) |
| phase | deg | nucleosome | (Satchwell, Drew, and Travers 1986) |
| Roll3 | deg | nucleosome | (Goodsell and Dickerson 1994) |
| MGW | A | DNAshapeR | (Chiu et al. 2016; Rohs et al. 2009) |
| ProT | deg | DNAshapeR | (Chiu et al. 2016; Rohs et al. 2009) |
| Roll | deg | DNAshapeR | (Chiu et al. 2016; Rohs et al. 2009) |
| HelT | deg | DNAshapeR | (Chiu et al. 2016; Rohs et al. 2009) |
| HRC | cleavage intensity | ORChID2 | (Bishop et al. 2011) |
| TIDD | no. of events | TIDD | (Zrimec and Lapanje 2015) |
| Tm | deg C | oligoprop | Matlab function |

**Table S1-4.** Alignment algorithm performance measures.

| Classification metric | Shorthand | Equation |
|---|---|---|
| **True positives** | TP | TP |
| **True negatives** | TN | TN |
| **False positives** | FP | FP |
| **False negatives** | FN | FN |
| **Sensitivity (Recall)** | TPR | TP/(TP+FN) |
| **Specificity** | TNR | TN/(TN+FP) |
| **Precision** | PPV | TP/(TP+FP) |
| **Accuracy** | Acc | (TP+TN)/(TP+FP+FN+TN) |
| **F1 score** | F1S | 2*TP/(2*TP+FP+FN) |
| **Matthews corr. coef.** | MCC | (TP*TN-FP*FN)/sqrt((TP+FP)*(TP+FN)*(TN+FP)*(TN+FN)) |

**Table S2-1.** Pearson correlation coefficients between sequence identities at all regions sizes. All p-values below 1e-16.

| **region_size 2** | **40** | **60** | **90** | **120** | **160** | **220** |
|---|---|---|---|---|---|---|
| **region_size 1** | | | | | | |
| **40** | 1 | 0.940277 | 0.916119 | 0.907978 | 0.889912 | 0.878428 |
| **60** | 0.940277 | 1 | 0.955234 | 0.940295 | 0.925025 | 0.911239 |
| **90** | 0.916119 | 0.955234 | 1 | 0.973206 | 0.959329 | 0.944463 |
| **120** | 0.907978 | 0.940295 | 0.973206 | 1 | 0.979166 | 0.964006 |
| **160** | 0.889912 | 0.925025 | 0.959329 | 0.979166 | 1 | 0.978883 |
| **220** | 0.878428 | 0.911239 | 0.944463 | 0.964006 | 0.978883 | 1 |

**Table S2-2.** Enrichment analysis of plasmid genomic location of oriT regions.

| Product type | Proportion | Fold change | Fisher's test p-value |
|---:|---:|---:|---:|
| **Other** | 0.501689 | 0.955218 | 6.41E-07 |
| **Hypothetical protein** | 0.197835 | 0.543307 | <1E-16 |
| **None** | 0.21256 | 4.569195 | 0.00E+00 |
| **Transposition** | 0.0466 | 1.323258 | 4.47E-10 |
| **Conjugation** | 0.025379 | 1.166235 | 1.06E-02 |
| **Integration** | 0.015938 | 2.225049 | <1E-16 |

**Table S2-3.** Counts of orit hits across mob groups and subgroups.

| mob | mob_subgroup | query_orits | orit_hits |
|---|---|---|---|
| **C** | **1** | 2 | 339 |
| | **2** | 2 | 12 |
| **F** | **11** | 6 | 82 |
| | **12** | 9 | 149 |
| | **13** | 1 | 2 |
| | **u** | 1 | 1512 |
| **P** | **11** | 2 | 74 |
| | **12** | 1 | 71 |
| | **13** | 1 | 35 |
| | **14** | 2 | 16 |
| | **2** | 1 | 2 |
| | **3** | 1 | 67 |
| | **4** | 5 | 189 |
| | **5** | 3 | 322 |
| | **6** | 3 | 13 |
| | **7** | 2 | 42 |
| **Q** | **1** | 7 | 140 |
| | **2** | 7 | 3264 |
| | **3** | 3 | 29 |
| | **u** | 10 | 3434 |
| **T** | **1** | 1 | 1 |
| **V** | **1** | 27 | 232 |
| | **2** | 1 | 1371 |
| | **3** | 2 | 1 |
| | **4** | 6 | 5 |
| | **u** | 6 | 93 |

**Table S3-1.** Top 30 sorted oriTs according to the degree of all and unique connections with other oriTs.

| node | Plasmid_name | Species | Mob | Mob subgroup | degree | degree_unique | num_plasmids | Connected Mob |
|---|---|---|---|---|---|---|---|---|
| 50 | pNL1 | *Novosphingobium aromaticivorans* | F | u | 18456 | 56 | 1512 | (P, V, Q, F, T, C) |
| 64 | BNC1 Plasmid 1 | *Mesorhizobium sp.* | Q | 2 | 22651 | 54 | 1772 | (P, V, Q, F, T, C) |
| 31 | pBBR1 | *Bordetella bronchiseptica* | V | 2 | 20875 | 50 | 1371 | (P, V, Q, F, T, C) |
| 72 | pDOJH10S | *Bifidobacterium longum* | Q | u | 24632 | 42 | 1617 | (P, V, Q, F, T, C) |
| 69 | pKJ50 | *Bifidobacterium longum* | Q | u | 15585 | 42 | 921 | (P, V, Q, F, T, C) |
| 24 | pTiC58 | *Agrobacterium tumefaciens* | Q | 2 | 13879 | 41 | 883 | (P, V, Q, F, T, C) |
| 13 | ColE1 | *Escherichia coli* | P | 5 | 3179 | 40 | 250 | (P, V, Q, F, T, C) |
| 71 | pMG160 | *Rhodobacter blasticus* | Q | u | 11863 | 38 | 698 | (P, V, Q, F, T, C) |
| 35 | CloDF13 | *Escherichia coli* | C | 1 | 6429 | 36 | 334 | (P, V, Q, F, T, C) |
| 21 | pSymB | *Sinorhizobium meliloti* | Q | 2 | 3535 | 33 | 192 | (P, V, Q, F, C) |
| 25 | pRetCFN42d | *Rhizobium etli* | Q | 2 | 3608 | 32 | 175 | (P, V, Q, F, C) |
| 20 | pNGR234a | *Sinorhizobium fredii* | Q | 2 | 3161 | 30 | 171 | (P, V, Q, F, C) |
| 27 | pTF1 | *Acidithiobacillus ferrooxidans* | Q | u | 2610 | 30 | 136 | (P, V, Q, F, C) |
| 108 | pWKS1 | *Paracoccus pantotrophus* | V | u | 1180 | 29 | 70 | (P, V, Q, F, C) |
| 60 | pIE1130 | *uncultured eubacterium* | Q | 1 | 1425 | 28 | 88 | (P, V, Q, F, C) |
| 22 | pSymA | *Sinorhizobium meliloti* | Q | 2 | 718 | 25 | 40 | (P, V, Q, F, C) |
| 51 | pRA2 | *Pseudomonas alcaligenes* | P | 13 | 650 | 24 | 35 | (P, V, Q, F, C) |
| 107 | pOM1 | *Oenococcus oeni* | V | u | 175 | 24 | 19 | (P, V, Q, F, C) |
| 23 | p42a | *Rhizobium etli* | Q | 2 | 600 | 23 | 31 | (P, V, Q, F, C) |

| | | | | | | | | |
|---|---|---|---|---|---|---|---|---|
| 29 | pDN1 | *Dichelobacter nodosus* | Q | u | 476 | 23 | 23 | (P, V, Q, F, C) |
| 61 | pIE1115 | *uncultured eubacterium* | Q | 1 | 360 | 23 | 24 | (P, V, Q, F, C) |
| 56 | pWQ799 | *Salmonella enterica* | P | 5 | 136 | 20 | 70 | (P, V, Q, F, C) |
| 7 | R64 | *Sinorhizobium fredii* | P | 12 | 100 | 18 | 71 | (P, V, Q, F) |
| 53 | pTF-FC2 | *Acidithiobacillus ferrooxidans* | P | 14 | 103 | 18 | 11 | (P, V, Q, F, C) |
| 5 | RP4/pTB11/ pBS228 | *Escherichia coli* | P | 11 | 219 | 17 | 30 | (P, V, Q, F, C) |
| 11 | pVT745 | *Aggregatibacter actinomycetemcomitans* | P | 4 | 95 | 17 | 184 | (P, V, Q, F) |
| 68 | pDOJH10L | *Bifidobacterium longum* | Q | u | 70 | 16 | 12 | (P, V, Q, F, C) |
| 73 | pSmeLPU88b | *Sinorhizobium meliloti* | Q | u | 107 | 16 | 9 | (P, V, Q, F, C) |
| 52 | pTC-F14 | *Acidithiobacillus caldus* | P | 14 | 53 | 15 | 5 | (P, V, Q, F, C) |
| 6 | R751/pTP6 | *Enterobacter aerogenes* | P | 11 | 302 | 15 | 44 | (P, V, Q, F, C) |

**Table S3-2.** Orit and relaxase typing results partitioned across mob groups, showing the number of unique plasmids, species, oriTs and connections.

| | MOB | C | F | H | P | Q | T | V |
|---|---|---|---|---|---|---|---|---|
| Relaxase | No. plasmids | 35 | 305 | 82 | 418 | 277 | NaN | 241 |
| | No. species | 20 | 92 | 38 | 170 | 112 | NaN | 88 |
| | No. oriT (MOB) | 35 | 306 | 83 | 423 | 284 | NaN | 246 |
| | No. conn.* | 35 | 306 | 83 | 423 | 284 | NaN | 246 |
| | Avg. conn. | 0.057143 | 0.045752 | 0.024096 | 0.236407 | 0.197183 | NaN | 0.479675 |
| | Avg. MOB conn.** | 0.028571 | 0.022876 | 0.012048 | 0.118203 | 0.03169 | NaN | 0.178862 |
| oriT | No. plasmids | 253 | 1047 | NaN | 694 | 1274 | 1 | 923 |
| | No. species | 137 | 304 | NaN | 217 | 398 | 1 | 322 |
| | No. oriT | 351 | 1745 | NaN | 831 | 6867 | 1 | 1702 |
| | No. conn.* | 351 | 1745 | NaN | 831 | 6867 | 1 | 1702 |
| | Avg. conn. | 485.891738 | 224.973066 | NaN | 136.815884 | 365.444008 | 650 | 301.79436 |
| | Avg. MOB conn.** | 3.014245 | 2.102006 | NaN | 1.33935 | 2.373671 | 5 | 2.277321 |
| Rel. increase | No. plasmids | 7.228571 | 3.432787 | NaN | 1.660287 | 4.599278 | NaN | 3.829876 |
| | No. species | 6.85 | 3.304348 | NaN | 1.276471 | 3.553571 | NaN | 3.659091 |
| | No. oriT | 10.028571 | 5.702614 | NaN | 1.964539 | 24.179577 | NaN | 6.918699 |
| | No. conn.* | 10.028571 | 5.702614 | NaN | 1.964539 | 24.179577 | NaN | 6.918699 |
| | Avg. conn. | 8503.105413 | 4917.26844 | NaN | 578.731191 | 1853.323181 | NaN | 629.164512 |
| | Avg. MOB conn.** | 105.498575 | 91.887679 | NaN | 11.330903 | 74.902513 | NaN | 12.732294 |

* length(n)*(length(n)-1)

** length(unique(n))-1

**Table S3-3.** Adjacency matrix of MOB connections across plasmids (note: symmetric across diagonal).

| Mob | C | F | P | Q | T | V |
|---|---|---|---|---|---|---|
| **C** | 122 | 742 | 195 | 4371 | 1 | 889 |
| **F** | 742 | 970 | 664 | 12637 | 4 | 2806 |
| **P** | 195 | 664 | 140 | 3118 | 1 | 743 |
| **Q** | 4371 | 12637 | 3118 | 35062 | 17 | 15081 |
| **T** | 1 | 4 | 1 | 17 | 0 | 2 |
| **V** | 889 | 2806 | 743 | 15081 | 2 | 1439 |

**Table S3-4.** Top 20 sorted connections between Mob subgroups.

| Mob subgroup 1 | Mob subgroup 2 | Num. connections |
|---|---|---|
| **Q2** | **Qu** | 15983 |
| **Qu** | **Qu** | 10223 |
| **Q2** | **V1** | 10223 |
| | **Fu** | 8972 |
| **Fu** | **Qu** | 8972 |
| **V2** | **Qu** | 7309 |
| **Q2** | **Vu** | 7309 |
| | **Q2** | 7268 |
| **P7** | **Qu** | 7268 |
| **T1** | **Qu** | 3534 |
| **Q2** | **V2** | 3534 |
| **C1** | **Qu** | 3274 |
| **Q2** | **C1** | 3274 |
| **V2** | **Q2** | 3095 |
| **P7** | **Vu** | 3095 |
| | **Fu** | 2916 |
| **Fu** | **Q2** | 2916 |
| **V2** | **Fu** | 2737 |
| **Fu** | **Vu** | 2737 |
| **T1** | **Vu** | 1220 |

**Table S4-1.** Species and genus counts across environmental habitats.

| Supertype | Type | Subtype | Species | Genus |
|---|---|---|---|---|
| **Animal** | **Animal** | **Animal** | 1453 | 1453 |
| **Freshwater** | **Freshwater** | **Freshwater** | 2814 | 2814 |
| **Human** | **Human_commensal** | **Human_Gut** | 393 | 254 |
| | | **Human_Oral** | 259 | 221 |
| | | **Human_Other tissues** | 234 | 234 |
| | | **Human_Respiratory** | 32 | 7 |
| | | **Human_Skin** | 19 | 2 |
| | | **Human_Unassigned** | 730 | 196 |
| | | **Human_Urogenital** | 10 | 3 |
| | **Human_pathogen** | **Human_Gut** | 60 | 47 |
| | | **Human_Oral** | 62 | 55 |
| | | **Human_Respiratory** | 39 | 35 |
| | | **Human_Skin** | 48 | 43 |
| | | **Human_Unassigned** | 55 | 49 |
| | | **Human_Urogenital** | 29 | 23 |
| **Industrial** | **Industrial** | **Industrial** | 1879 | 1879 |
| **Interface** | **Interface** | **Interface** | 936 | 936 |
| **Plant** | **Plant** | **Plant** | 307 | 307 |
| **Saline** | **Saline** | **Saline** | 2084 | 2084 |
| **Soil** | **Soil** | **Soil** | 2707 | 2707 |
| **Thermal** | **Thermal** | **Thermal** | 439 | 439 |

**Table S4-2.** Used habitat groupings corresponding to MicroDB.

| MicroDB supertype | MicroDB type | MicroDB subtype | Species | Genus | Used supertype |
|---|---|---|---|---|---|
| **Artificial** | **Compost** | **unclassified** | 248 | 248 | Industrial |
| | **Foods and drinks** | **unclassified** | 203 | 203 | Industrial |
| | **Water treatment** | **unclassified** | 685 | 685 | Industrial |
| | **unclassified** | **unclassified** | 33 | 33 | Industrial |
| **Host-associated** | **Gut** | **Cattle** | 219 | 219 | Animal |
| | | **Human** | 453 | 301 | Human |
| | | **Insect** | 78 | 78 | Animal |
| | | **Mouse** | 50 | 50 | Animal |
| | | **Other** | 319 | 319 | Animal |
| | | **unclassified** | 261 | 261 | Animal |
| | **Marine host-associated** | **unclassified** | 227 | 227 | Animal |
| | **Oral** | **Human** | 321 | 276 | Human |
| | **Other tissues** | **Human** | 234 | 234 | Human |
| | | **Other** | 299 | 299 | Animal |
| | **Plants** | **unclassified** | 307 | 307 | Plant |
| | **Respiratory** | **Human** | 71 | 42 | Human |
| | **Skin** | **Human** | 67 | 45 | Human |
| | **Unassigned** | **Human** | 785 | 245 | Human |
| | **Urogenital** | **Human** | 39 | 26 | Human |
| | **unclassified** | **unclassified** | 286 | 286 | / |
| **Hypothermal** | **Marine** | **unclassified** | 80 | 80 | Saline |
| | **Non-marine** | **unclassified** | 334 | 334 | Freshwater |
| **Non-saline** | **Freshwaters** | **Aquifer** | 296 | 296 | Freshwater |
| | | **Drinking water** | 254 | 254 | Freshwater |
| | | **Groundwater** | 338 | 338 | Freshwater |
| | | **Interface** | 48 | 48 | Freshwater |
| | | **Lake** | 492 | 492 | Freshwater |
| | | **Lakes** | 62 | 62 | Freshwater |
| | | **River** | 296 | 296 | Freshwater |
| | | **Sediment** | 351 | 351 | Freshwater |
| | | **unclassified** | 343 | 343 | Freshwater |

| | | | | | |
|---|---|---|---|---|---|
| | **Interface** | **unclassified** | 195 | 195 | Interface |
| | **Soils** | **Agricultural** | 368 | 368 | Soil |
| | | **Arid** | 165 | 165 | Soil |
| | | **Caves/Rocks/Mines** | 358 | 358 | Soil |
| | | **Forest** | 193 | 193 | Soil |
| | | **Grassland** | 149 | 149 | Soil |
| | | **Other** | 112 | 112 | Soil |
| | | **Wetland** | 308 | 308 | Soil |
| | | **unclassified** | 1054 | 1054 | Soil |
| | **unclassified** | **unclassified** | 209 | 209 | Interface |
| **Other** | **Aerial** | **unclassified** | 532 | 532 | Interface |
| | **Oil** | **unclassified** | 366 | 366 | Industrial |
| | **Other** | **Industrial** | 344 | 344 | Industrial |
| | **unclassified** | **unclassified** | 2 | 2 | / |
| **Saline** | **Hypersaline** | **Lakes** | 164 | 164 | Saline |
| | | **Saltern** | 16 | 16 | Saline |
| | | **unclassified** | 96 | 96 | Saline |
| | **Interfaces** | **unclassified** | 99 | 99 | Saline |
| | **Marine** | **Sediment** | 393 | 393 | Saline |
| | | **Water** | 573 | 573 | Saline |
| | | **unclassified** | 329 | 329 | Saline |
| | **Soils** | **unclassified** | 225 | 225 | Saline |
| | **unclassified** | **unclassified** | 109 | 109 | Saline |
| **Thermal** | **Other** | **unclassified** | 34 | 34 | Thermal |
| | **Springs** | **unclassified** | 240 | 240 | Thermal |
| | **Vents** | **Other** | 1 | 1 | Thermal |
| | | **unclassified** | 157 | 157 | Thermal |
| | **unclassified** | **unclassified** | 7 | 7 | Thermal |
| **unclassified** | **unclassified** | **unclassified** | 1195 | 1195 | / |

**Table S5-1.** Author contributions as defined by the CRediT taxonomy (https://casrai.org/credit/).

| Author | Conceptualization | Data curation | Formal Analysis | Funding acquisition | Investigation | Methodology | Project administration | Resources | Software | Supervision | Validation | Visualization | Writing – original draft | Writing – review & editing |
|---|---|---|---|---|---|---|---|---|---|---|---|---|---|---|
| JZ | x | x | x | x | x | x | x | x | x | x | x | x | x | x |

### Supplementary references

Aida, Misako. 1988. “An Ab Initio Molecular Orbital Study on the Sequence-Dependency of DNA Conformation: An Evaluation of Intra- and Inter-Strand Stacking Interaction Energy.” *Journal of Theoretical Biology*. https://doi.org/10.1016/s0022-5193(88)80032-8.

Bishop, Eric P., Remo Rohs, Stephen C. J. Parker, Sean M. West, Peng Liu, Richard S. Mann, Barry Honig, and Thomas D. Tullius. 2011. “A Map of Minor Groove Shape and Electrostatic Potential from Hydroxyl Radical Cleavage Patterns of DNA.” *ACS Chemical Biology* 6 (12): 1314–20.

Bolshoy, A., and P. McNamara. 1991. “Curved DNA without AA: Experimental Estimation of All 16 DNA Wedge Angles.” *Proceedings of the*. https://www.pnas.org/content/88/6/2312.short.

Breslauer, K. J., R. Frank, H. Blöcker, and L. A. Marky. 1986. “Predicting DNA Duplex Stability from the Base Sequence.” *Proceedings of the National Academy of Sciences of the United States of America* 83 (11): 3746–50.

Brukner, I., R. Sánchez, D. Suck, and S. Pongor. 1995. “Trinucleotide Models for DNA Bending Propensity: Comparison of Models Based on DNaseI Digestion and Nucleosome Packaging Data.” *Journal of Biomolecular Structure & Dynamics* 13 (2): 309–17.

Brukner, Ivan, Roberto Sanchez, Dietrich Suck, and Sandor Pongor. 1995. “Sequence-Dependent Bending Propensity of DNA as Revealed by DNase I: Parameters for Trinucleotides.” *The EMBO Journal* 14 (8): 1812–18.

Chiu, Tsu-Pei, Federico Comoglio, Tianyin Zhou, Lin Yang, Renato Paro, and Remo Rohs. 2016. “DNAshapeR: An R/Bioconductor Package for DNA Shape Prediction and Feature Encoding.” *Bioinformatics* 32 (8): 1211–13.

Dewhirst, Floyd E., Tuste Chen, Jacques Izard, Bruce J. Paster, Anne C. R. Tanner, Wen-Han Yu, Abirami Lakshmanan, and William G. Wade. 2010. “The Human Oral Microbiome.” *Journal of Bacteriology* 192 (19): 5002–17.

Escapa, Isabel F., Tsute Chen, Yanmei Huang, Prasad Gajare, Floyd E. Dewhirst, and Katherine P. Lemon. 2018. “New Insights into Human Nostril Microbiome from the Expanded Human Oral Microbiome Database (eHOMD): A Resource for the Microbiome of the Human Aerodigestive Tract.” *mSystems* 3 (6). https://doi.org/10.1128/mSystems.00187-18.

Forster, Samuel C., Hilary P. Browne, Nitin Kumar, Martin Hunt, Hubert Denise, Alex Mitchell, Robert D. Finn, and Trevor D. Lawley. 2016. “HPMCD: The Database of Human Microbial Communities from Metagenomic Datasets and Microbial Reference Genomes.” *Nucleic Acids Research* 44 (D1): D604–9.

Gartenberg, M. R., and D. M. Crothers. 1988. “DNA Sequence Determinants of CAP-Induced Bending and Protein Binding Affinity.” *Nature* 333 (6176): 824–29.

Geggier, Stephanie, and Alexander Vologodskii. 2010. “Sequence Dependence of DNA Bending Rigidity.” *Proceedings of the National Academy of Sciences of the United States of America* 107 (35): 15421–26.

Goodsell, D. S., and R. E. Dickerson. 1994. “Bending and Curvature Calculations in B-DNA.” *Nucleic Acids Research* 22 (24): 5497–5503.

Gorin, A. A., V. B. Zhurkin, and W. K. Olson. 1995. “B-DNA Twisting Correlates with Base-Pair Morphology.” *Journal of Molecular Biology* 247 (1): 34–48.

Gotoh, Osamu, and Yusaku Tagashira. 1981. “Stabilities of Nearest-Neighbor Doublets in Double-Helical DNA Determined by Fitting Calculated Melting Profiles to Observed Profiles.” *Biopolymers: Original Research on Biomolecules* 20 (5): 1033–42.

Hartmann, B., B. Malfoy, and R. Lavery. 1989. “Theoretical Prediction of Base Sequence Effects in DNA. Experimental Reactivity of Z-DNA and B-Z Transition Enthalpies.” *Journal of Molecular Biology* 207 (2): 433–44.

Ho, Pui S., Michael J. Ellison, Gary J. Quigley, and Alexander Rich. 1986. “A Computer Aided Thermodynamic Approach for Predicting the Formation of Z-DNA in Naturally Occurring Sequences.” *The EMBO Journal* 5 (10): 2737–44.

Human Microbiome Project Consortium. 2012. “Structure, Function and Diversity of the Healthy Human Microbiome.” *Nature* 486 (7402): 207–14.

Kabsch, W., C. Sander, and E. N. Trifonov. 1982. “The Ten Helical Twist Angles of B-DNA.” *Nucleic Acids Research*. https://doi.org/10.1093/nar/10.3.1097.

Karas, H., R. Knüppel, W. Schulz, H. Sklenar, and E. Wingender. 1996. “Combining Structural Analysis of DNA with Search Routines for the Detection of Transcription Regulatory Elements.” *Computer Applications in the Biosciences: CABIOS* 12 (5): 441–46.

Kulkarni, Mandar, and Arnab Mukherjee. 2013. “Sequence Dependent Free Energy Profiles of Localized B- to A-Form Transition of DNA in Water.” *The Journal of Chemical Physics* 139 (15): 155102.

Li, Xiaobin, Yingzhou Xie, Meng Liu, Cui Tai, Jingyong Sun, Zixin Deng, and Hong-Yu Ou. 2018. “oriTfinder: A Web-Based Tool for the Identification of Origin of Transfers in DNA Sequences of Bacterial Mobile Genetic Elements.” *Nucleic Acids Research* 46 (W1): W229–34.

Lloyd-Price, Jason, Anup Mahurkar, Gholamali Rahnavard, Jonathan Crabtree, Joshua Orvis, A. Brantley Hall, Arthur Brady, et al. 2017. “Strains, Functions and Dynamics in the Expanded Human Microbiome Project.” *Nature* 550 (7674): 61–66.

Olson, Wilma K., Andrey A. Gorin, Xiang-Jun Lu, Lynette M. Hock, and Victor B. Zhurkin. 1998. “DNA Sequence-Dependent Deformability Deduced from protein–DNA Crystal Complexes.” *Proceedings of the National Academy of Sciences of the United States of America* 95 (19): 11163–68.

Packer, M. J., M. P. Dauncey, and C. A. Hunter. 2000. “Sequence-Dependent DNA Structure: Dinucleotide Conformational Maps.” *Journal of Molecular Biology* 295 (1): 71–83.

Perez, Alberto, Agnes Noy, Filip Lankas, F. Javier Luque, and Modesto Orozco. 2004. “The Relative Flexibility of B-DNA and A-RNA Duplexes: Database Analysis.” *Nucleic Acids Research* 32 (20): 6144–51.

Pignatelli, Miguel, Andrés Moya, and Javier Tamames. 2009. “EnvDB, a Database for Describing the Environmental Distribution of Prokaryotic Taxa.” *Environmental Microbiology Reports* 1 (3): 191–97.

Protozanova, Ekaterina, Peter Yakovchuk, and Maxim D. Frank-Kamenetskii. 2004.

"Stacked–Unstacked Equilibrium at the Nick Site of DNA." *Journal of Molecular Biology*. https://doi.org/10.1016/j.jmb.2004.07.075.

Rohs, Remo, Sean M. West, Alona Sosinsky, Peng Liu, Richard S. Mann, and Barry Honig. 2009. "The Role of DNA Shape in protein–DNA Recognition." *Nature* 461 (7268): 1248–53.

SantaLucia, J., Jr. 1998. "A Unified View of Polymer, Dumbbell, and Oligonucleotide DNA Nearest-Neighbor Thermodynamics." *Proceedings of the National Academy of Sciences of the United States of America* 95 (4): 1460–65.

Satchwell, S. C., H. R. Drew, and A. A. Travers. 1986. "Sequence Periodicities in Chicken Nucleosome Core DNA." *Journal of Molecular Biology* 191 (4): 659–75.

Seyed-Allaei, Hamed, Ginestra Bianconi, and Matteo Marsili. 2006. "Scale-Free Networks with an Exponent Less than Two." *Physical Review. E, Statistical, Nonlinear, and Soft Matter Physics* 73 (4 Pt 2): 046113.

Sivolob, A. V., and S. N. Khrapunov. 1995. "Translational Positioning of Nucleosomes on DNA: The Role of Sequence-Dependent Isotropic DNA Bending Stiffness." *Journal of Molecular Biology* 247 (5): 918–31.

Zrimec, Jan, and Ales Lapanje. 2015. "Fast Prediction of DNA Melting Bubbles Using DNA Thermodynamic Stability." *IEEE/ACM Transactions on Computational Biology and Bioinformatics / IEEE, ACM* 12 (5): 1137–45.

Zrimec, Jan, and Aleš Lapanje. 2018. "DNA Structure at the Plasmid Origin-of-Transfer Indicates Its Potential Transfer Range." *Scientific Reports*. https://doi.org/10.1038/s41598-018-20157-y.
